# Altered Hippocampal-Prefrontal Neural Dynamics in Mouse Models of Down Syndrome

**DOI:** 10.1101/644849

**Authors:** Pishan Chang, Daniel Bush, Stephanie Schorge, Mark Good, Tara Canonica, Nathanael Shing, Suzanna Noy, Frances K. Wiseman, Neil Burgess, Victor L.J. Tybulewicz, Matthew C. Walker, Elizabeth M.C. Fisher

## Abstract

Altered neural dynamics in medial prefrontal cortex (mPFC) and hippocampus may contribute to cognitive impairments in the complex chromosomal disorder, Down Syndrome (DS). Here, we demonstrate non-overlapping behavioural differences associated with distinct abnormalities in hippocampal and mPFC electrophysiology during a canonical spatial memory task in three partially trisomic mouse models of DS (Dp1Tyb, Dp10Yey, Dp17Yey) that together cover all regions of homology with human chromosome 21 (Hsa21). Dp1Tyb mice showed slower decision-making (unrelated to the gene dose of *DYRK1A*, which has been implicated in DS cognitive dysfunction) and altered theta dynamics (reduced frequency, increased hippocampal-mPFC coherence, increased modulation of hippocampal high gamma); Dp10Yey mice showed impaired alternation performance and reduced theta modulation of hippocampal low gamma; while Dp17Yey mice were no different from wildtype mice. These results link specific hippocampal and mPFC circuit dysfunctions to cognitive deficits in DS models and, importantly, map them to discrete regions of Hsa21.

## Introduction

Down syndrome (DS) is a complex cognitive disorder arising from trisomy of human chromosome 21 (Hsa21) with an incidence of ∼1 in 800 live births worldwide^1^. The current global population of people with DS is estimated at 6 million^2^ and prevalence is rising, primarily due to an increase in maternal age (a major risk factor for DS) and increased life expectancy in people with DS resulting from reduced infant mortality rates and improved access to healthcare^3–5^. DS is characterised by intellectual disability^6, 7^ and prominent impairments in planning, decision-making, and memory function^7–12^ which likely arise from functional abnormalities of hippocampus and medial prefrontal cortex (mPFC)^6, 8, 9, 13–16^. Increased dosage of single genes in Hsa21, such as *DYRK1A*, have been proposed to account for many of the alterations in neural development and abnormal phenotypes associated with DS, and thus to be targets for therapy development^17^.

Activity in the hippocampus and mPFC can be characterised by oscillations in the theta and gamma bands. Hippocampal theta oscillations are associated with translational movement^18, 19^ and mnemonic function^20–22^ across species, and can modulate synaptic plasticity^23^. Moreover, hippocampal theta modulates the amplitude of concomitant gamma oscillations both locally and across the neocortex^24–26^, and task-related increases in phase-amplitude coupling are associated with successful memory encoding^27^. In humans, theta oscillations in mPFC are observed during working memory maintenance^28, 29^ and long-term memory retrieval^30, 31^, while increases in theta coherence between hippocampus and mPFC are associated with planning and decision-making across species^32–36^.

To further elucidate the neural mechanisms underlying cognitive deficits associated with DS, we studied three chromosome engineered mouse models that each exhibit trisomy for one region of orthology with human chromosome Hsa21 – referred to here as the Dp1Tyb, Dp10Yey and Dp17Yey strains (full nomenclature given in Materials and Methods)^37, 38^. In combination, these three mouse strains are triplicated for almost all the genes on Hsa21. We hypothesised that these trisomic mice might exhibit distinct cognitive impairments, corresponding to distinct alterations in oscillatory activity patterns within hippocampus and mPFC^15, 16^. Hence, we carried out simultaneous local field potential (LFP) recordings from those regions while mice performed a canonical spatial alternation task which, importantly, can dissociate mnemonic function (i.e. alternation success)^39–41^ from planning and decision-making processes (i.e. trial latency)^42–45^.

Here, we demonstrate that distinct behavioural impairments associated with DS are exhibited by animals with different regions of homology and, crucially, that these impairments are associated with distinct alterations in neural dynamics across hippocampus and mPFC. Moreover, reducing the ‘dose’ of *Dyrk1a* – a gene which has been suggested to be critically important for neural function in DS^46–51^ – was not sufficient to rescue the observed differences in behaviour, supporting the concept that cognitive impairments in DS do not necessarily map to single genes. By taking an unbiased approach to the gene content of these partially trisomic mice, and by combining behavioural and electrophysiological methodologies, we have therefore identified critical circuit dysfunction in DS models that paves the way for future determination of key dosage-sensitive gene combinations underlying cognitive phenotypes in this complex chromosomal disorder.

## Results

### Impaired Spatial Memory in Dp10Yey Mice and Decision-Making in Dp1Tyb Mice

Impairments in planning, decision making, and memory function have a significant impact on the lives of people with DS. In order to dissect the mechanisms underlying these cognitive deficits, we studied three mouse lines that are triplicated for the three mouse chromosome regions syntenic to Hsa21. The Dp1Tyb mouse strain has a 23Mb duplication of the Hsa21-syntenic region of Mmu16 which contains 148 coding genes with orthologues on Hsa21^37^; the Dp10Yey strain is duplicated for the Hsa21-syntenic region of Mmu10, which encodes 39 Hsa21 protein coding genes; and the Dp17Yey line is duplicated for the Hsa21-syntenic region of Mmu17 which encodes 19 protein coding genes^38^. Together, these mice make up a ‘mapping panel’, such that phenotypes found in any one strain are likely to arise from having an additional (i.e. third) copy of the specific Hsa21 orthologues within that strain.

We began by comparing cognitive function in male Dp1Tyb, Dp17Yey and Dp10Yey mice at 3 months of age with age- and sex-matched WT littermate control cohorts using a canonical spatial alternation task (Figure 1a-c; see Supplementary Figure 1a,b for further details and Supplementary Table 1 for trial and animal numbers). Importantly, this task can assay both mnemonic function (by examining the propensity to spontaneously alternate between goal arms on successive trials) and decision making (by examining the time taken to choose and enter a goal arm). Intriguingly, we found that distinct functional deficits were exhibited by each mutant mouse strain, suggesting that trisomy of discrete Hsa21 orthologues can have divergent effects on cognitive function.

**Figure 1:**
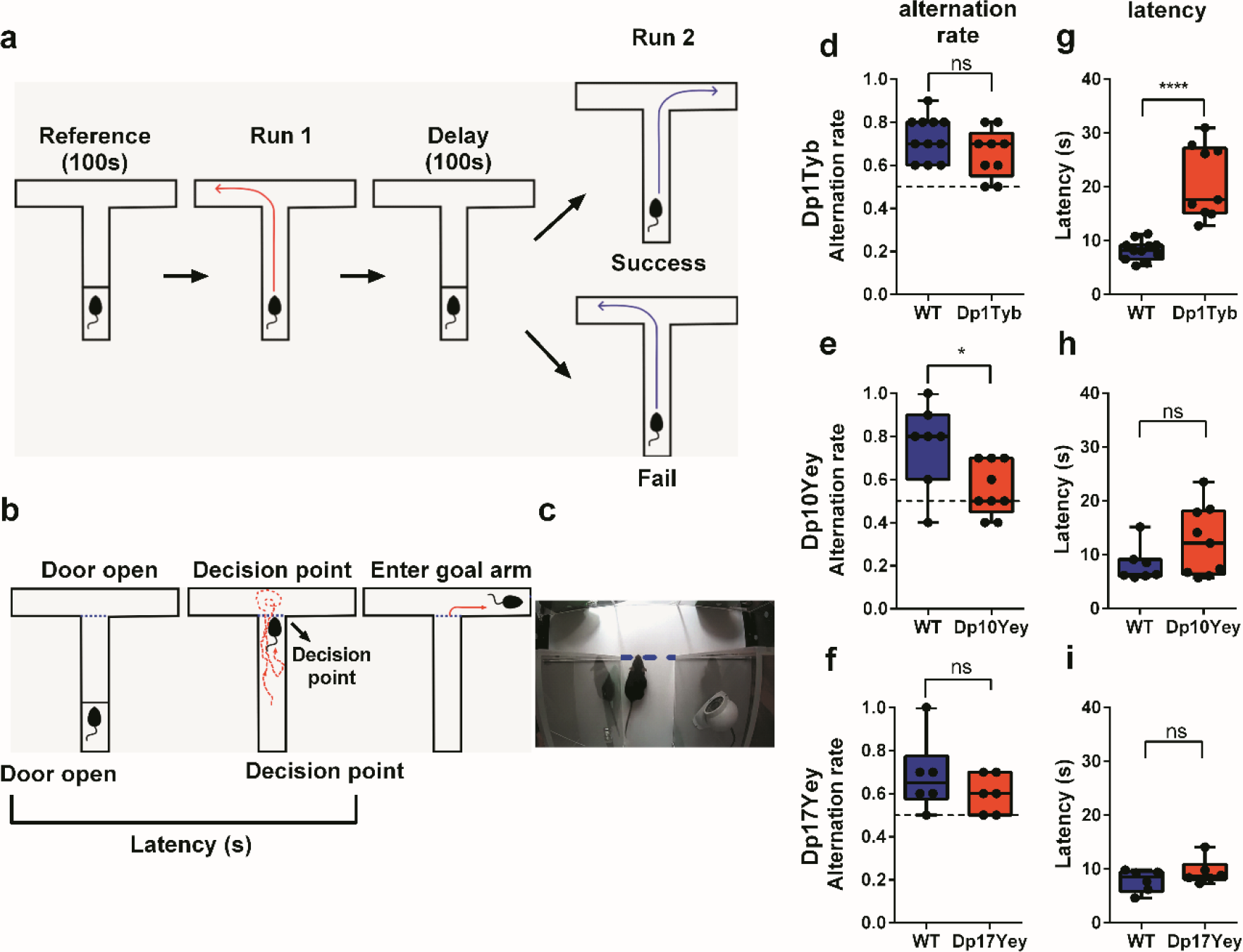
Spatial Alternation Rate and Trial Latency in Mouse Models of DS. **(a)** Schematic experimental procedure (see Supplementary Figure 1a,b for further details). **(b)** Schematic method for computing trial latency: the time between raising the door to release the animal from the start area, to the point at which the mouse’s nose crosses the ‘decision point’ (blue dashed line) before entering a goal arm. **(c)** Example of one animal reaching the decision point. **(d-f)** Alternation rate and **(g-i)** trial latency for each DS mouse model compared to their wild-type (WT) control group, showing: (e) significant differences in alternation rate for Dp10Yey vs WT mice (t(14)=2.48, p<0.05); and (g) significant differences in trial latency for Dp1Tyb vs WT mice (t(18)=5.97, p<0.001); but no differences in either measure for Dp17Yey vs WT mice. Chance alternation rate is shown as a black dotted line. Data are presented as box-whisker plots indicating the median, 25th and 75th percentiles, minimum and maximum values, with data for individual mice superimposed. Please refer to Supplementary Table 1 for trial and animal numbers, and Supplementary Table 2 for full details of all statistical analyses.

First, we found that Dp10Yey mice exhibited alternation rates that were significantly lower than their WT littermates and did not differ from chance (Figure 1e). In contrast, alternation rates in Dp1Tyb and Dp17Yey mice did not differ from those of WT mice and were significantly above chance in both strains (Figure 1d,f), with no difference in alternation rates between WT cohorts (Supplementary Figure 2a). These results suggest that Dp10Yey mice have impaired memory function, while Dp1Tyb and Dp17Yey mice are spared.

Second, we examined trial latencies, defined as the time taken to make a final crossing of the decision point prior to turning into the goal arm (see Materials and Methods). We found that these were significantly greater in Dp1Tyb mice compared to their WT littermates (Figure 1g); while no differences were observed between Dp10Yey or Dp17Yey mice and their respective control groups (Figure 1h,i) or between any of the WT cohorts (Supplementary Figure 2b). Importantly, the increased trial latency exhibited by Dp1Tyb mice could not simply be accounted for by motor impairments, independent of decision making processes, as we found no differences in average running speed between any mutant mouse group and their WT littermates. Conversely, Dp1Tyb mice spent a significantly greater proportion of each trial immobile, prior to making a decision, with no differences between either of the other mutant mouse strains and their control groups (Supplementary Figure 3). In sum, these results suggest that decision-making processes are disrupted in Dp1Tyb mice, despite relatively intact mnemonic function, while Dp10Yey and Dp17Yey mice are spared.

Finally, previous studies of transgenic mouse models of DS have led to the proposal that the overexpression of *Dyrk1a* (and thus an increased dosage of the DYRK1A protein) makes a critical contribution to neurological and behavioural abnormalities by shifting the excitation/inhibition balance towards inhibition, for example^16, 52^. The *Dyrk1a* gene maps to the Mmu16 region of Hsa21 and so is duplicated within the Dp1Tyb strain. To assess the behavioural consequences of altering the copy number of *Dyrk1a* in Dp1Tyb mice, we crossed Dp1Tyb animals with mice carrying a disrupted *Dyrk1a* gene to generate Dp1Tyb*Dyrk1aKO mice that are still duplicated for 147 Hsa21-orthologous coding genes on Mmu16, but have only 2 functional copies of *Dyrk1a*. Interestingly, these Dp1Tyb*Dyrk1aKO mice exhibited both a similar alternation rate to Dp1Tyb mice and a similarly prolonged decision-making (latency) phenotype (Supplementary Figure 4). Thus, reduction of the *Dyrk1a* copy number from three to two did not rescue the increased trial latency exhibited by Dp1Tyb mice. This finding indicates that triplication of *Dyrk1a* is not necessary to produce the decision-making deficit in Dp1Tyb mice, which must therefore arise from other gene(s) in this region of Hsa21 homology.

### Reduced Theta Frequency in Dp1Tyb Mice

Successful memory encoding and retrieval are associated with increased theta power in both hippocampus^20–22^ and mPFC^28–31^ across species. Furthermore, a reduction in hippocampal theta frequency has been directly linked to impaired spatial memory performance in a rodent model of temporal lobe epilepsy^53^. Hence, we next analysed LFP recordings from hippocampus and mPFC during spatial alternation in the T-maze (see Supplementary Figure 5 for details of electrode placement). Initially, we focused our analyses on a 10s window centred on the time at which animals crossed the decision point, and which would incorporate periods of memory encoding and retrieval from the sample and choice runs, respectively (see Materials and Methods for further details).

As expected, average power spectra from the mPFC (Figure 2a-c), and hippocampus (Figure 2d-f) across all animals showed a prominent peak in the 6-12Hz theta band during this period. Interestingly, although theta power did not differ between mouse lines, we found that theta frequency in both the mPFC (Figure 2a) and hippocampus (Figure 2d) was consistently lower in Dp1Tyb mice compared to WT controls. In contrast, no difference in theta frequency was observed in either region in Dp10Yey or Dp17Yey mice compared to their control cohorts (Figure 2b,c,e,f), or between WT cohorts (Supplementary Figure 2c,d).

**Figure 2:**
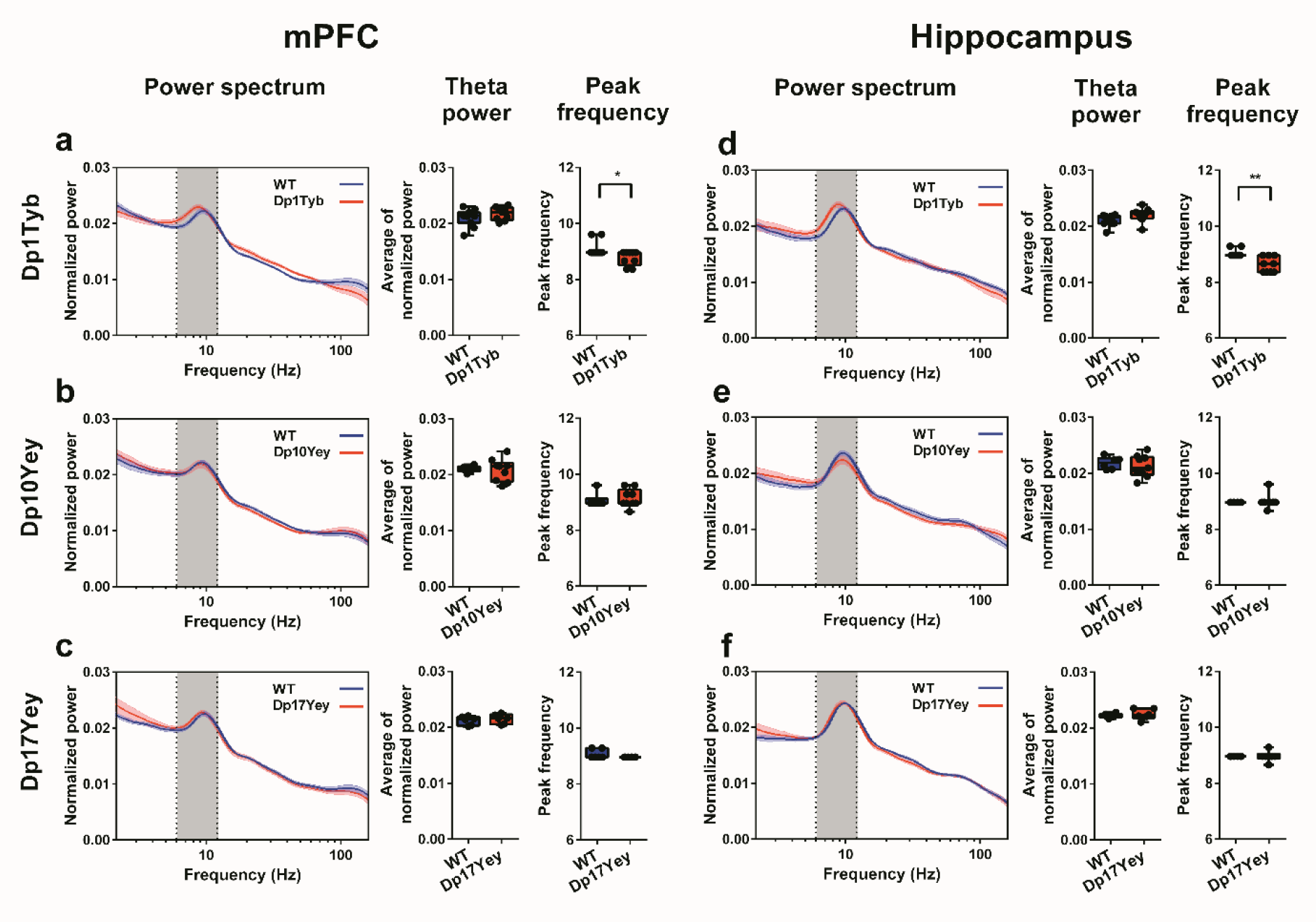
Theta Oscillations in mPFC and Hippocampus during Spontaneous Alternation. Power spectra, mean theta power and peak theta frequency in **(a-c)** mPFC and **(d-f)** hippocampus for **(a, d)** Dp1Tyb and WT; **(b, e)** Dp10Yey and WT; **(c, f)** Dp17Yey and WT animals during spontaneous alternation in the T-maze. Grey rectangles indicate the 6-12Hz theta band. There are no differences in theta power between mutant mice and WT groups in either mPFC or hippocampus. However, peak theta frequency in Dp1Tyb animals is significantly lower than WT in both: **(a)** mPFC (Dp1Tyb: 8.76±0.26Hz; WT: 9.08±0.26Hz; Mann–Whitney U=22.5, p<0.05); and **(d)** hippocampus (Dp1Tyb: 8.63±0.28Hz; WT: 9.02±0.13Hz; Mann–Whitney U=13.5, p<0.005), but no different in the other mutant mouse groups compared to their control populations (Mann–Whitney U test, all p>0.4). Data are presented as box-whisker plots indicating the median, 25th and 75th percentiles, minimum and maximum values, with data for individual mice superimposed. Please refer to Supplementary Table 2 for full details of all statistical analyses.

To establish whether the reduction in theta frequency observed in Dp1Tyb mice arose simply from the increased time that those animals spent immobile, we subsequently restricted our analysis to periods of movement only (see Materials and Methods). Consistent with the results above, theta frequency in both the hippocampus and mPFC of Dp1Tyb mice was still significantly lower than WT controls when periods of immobility were excluded. Moreover, this resulted from a reduction in the intercept, but not the slope, of the running speed v theta frequency relationship in both regions (Supplementary Figure 6a-d). In sum, these data suggest that Dp1Tyb mice, which exhibit slower decision making, also show a general slowing of theta band oscillations across hippocampal and medial prefrontal regions during spatial alternation, independent of running speed.

### Altered Hippocampal Phase-Amplitude Coupling in Dp1Tyb and Dp10Yey Mice

Coherence between the phase of theta oscillations and the amplitude of concurrent gamma band oscillations is prevalent in the rodent hippocampus^54, 55^ and across human neocortex^25^. In addition, theta-gamma phase-amplitude coupling (PAC) has been implicated in successful memory function^27, 56^. Hence, we asked whether the three DS mouse lines exhibited abnormal PAC that might be associated with the observed differences in behaviour. Average cross-frequency coherence images across all animals revealed two distinct PAC peaks in the hippocampal LFP: one between 6-12Hz theta and 60-120Hz ‘low gamma’ (LG) oscillations; and another between 6-12Hz theta and 140-160Hz ‘high gamma’ (HG) oscillations (Supplementary Figure 7a), while theta phase modulation of LG or HG amplitude was entirely absent in mPFC (Supplementary Figure 7b).

Interestingly, subsequent analyses indicated that the magnitude of hippocampal PAC in each pair of frequency bands also differed between specific DS models and WT controls. First, we found that theta-HG PAC was significantly increased in the Dp1Tyb group – which exhibited slowed decision making – relative to WT controls, but not in any other mouse strain (Figure 3a). Secondly, we found that theta-LG PAC was significantly reduced in the Dp10Yey group – which showed impaired spatial alternation – relative to WT controls, but not in any other strain (Figure 3b). Importantly, there was no alteration in hippocampal PAC across any pair of frequency bands in Dp17Yey animals, which also exhibit no differences in behaviour compared to their WT control group (Figure 3c), and no differences in theta-LG or theta-HG PAC between WT cohorts (Supplementary Figure 2e, f). In addition, we found no evidence for a difference in LG or HG power between mutant mice and their WT controls (Supplementary Figure 8).

**Figure 3:**
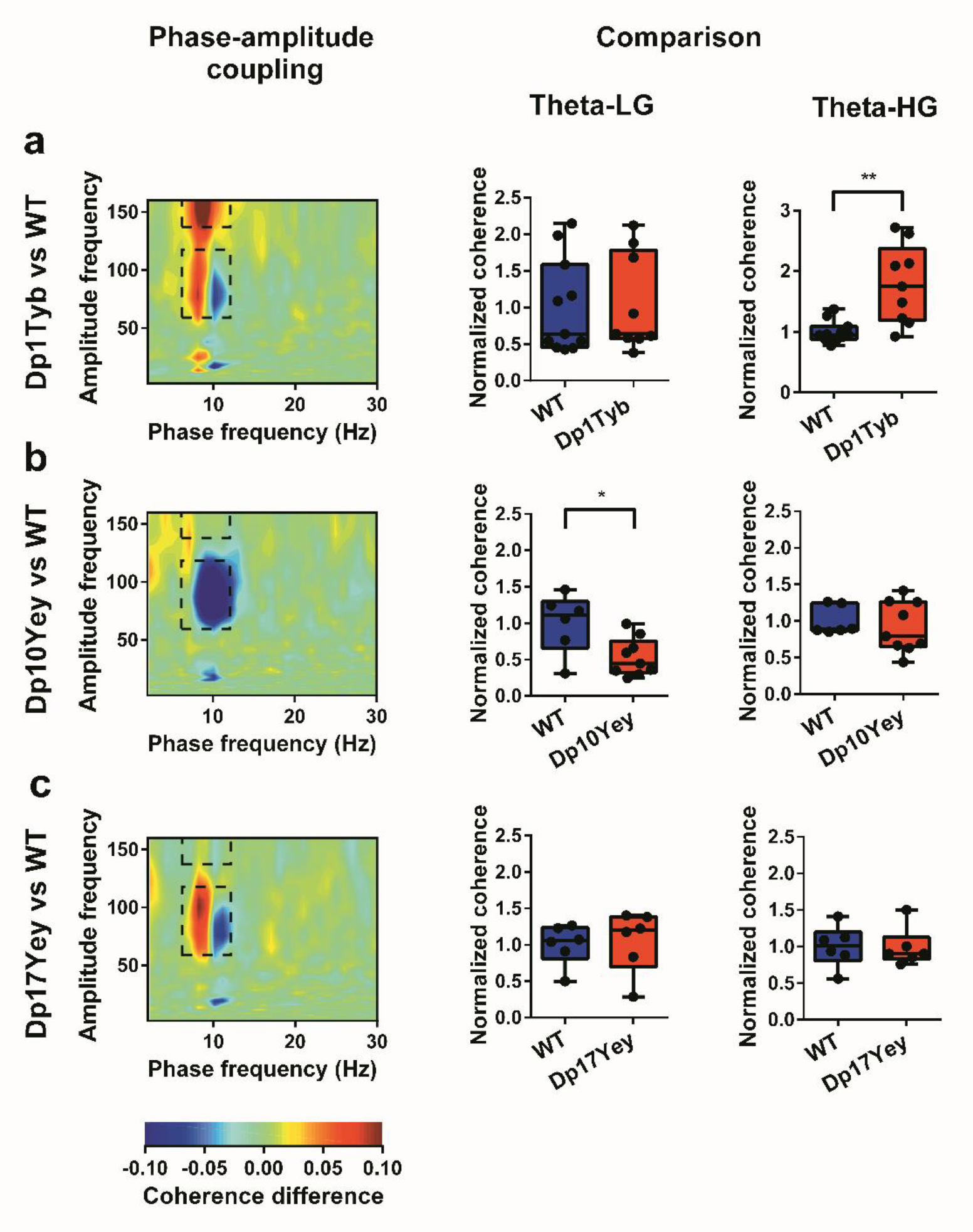
Hippocampal Phase-amplitude Coupling during Spontaneous Alternation. **(a-c)** Left: Comodulograms showing differences in hippocampal phase-amplitude coupling between each mutant mouse group and WT, with warm colours indicating stronger coupling in DS mice. These illustrate two prominent peaks – one between 6-12Hz theta phase and 60-120Hz ‘low gamma’ (LG) amplitude, and another between 6-12Hz theta phase and 140-160Hz ‘high gamma’ (HG) amplitude (black dashed rectangles; see Supplementary Figure 7 for further details). Right: Theta-LG and theta-HG cross-frequency coherence values, normalized by the mean value in the corresponding WT control cohort to facilitate comparison. **(a)** Dp1Tyb mice show no difference in theta-LG coupling, but significantly greater theta-HG coupling, compared to WT (Mann–Whitney U=11.0, p<0.01). **(b)** Conversely, Dp10Yey mice show significantly lower theta-LG coupling (Mann–Whitney U=8.0, p<0.05), but no difference in theta-HG coupling, compared to WT. **(c)** Finally, Dp17Yey mice show no difference in either theta-LG or theta-HG compared to WT. Data are presented as box-whisker plots indicating the median, 25th and 75th percentiles, minimum and maximum values, with data for individual mice superimposed. Please refer to Supplementary Table 2 for full details of all statistical analyses.

To confirm that the increased theta-HG PAC observed in Dp1Tyb mice did not arise from the observed differences in movement statistics, we subsequently removed any effect of average time spent immobile on average theta-HG PAC values across animals in both mutant and control groups by linear regression, and then compared the residual values between groups. This analysis confirmed that the increased theta-HG PAC in hippocampus exhibited by Dp1Tyb mice was independent of differences in movement statistics (Supplementary Figure 6e,f). In sum, these data distinguish changes in hippocampal theta phase modulation of local high (Dp1Tyb) and low (Dp10Yey) gamma amplitude in a manner that can be associated with increased trial latency and impaired spatial alternation, respectively.

### Increased Hippocampal-mPFC Theta Coherence in Dp1Tyb Mice

Planning, decision-making, memory encoding and retrieval processes are each associated with increased functional connectivity between the hippocampus and mPFC in both rodents^32–34, 57^ and humans^31, 36^. Interestingly, abnormalities in functional connectivity have also been implicated in various neurodevelopmental disorders, including DS^15, 58^. Hence, we next examined theta and gamma band coherence between hippocampus and mPFC, with the hypothesis that differences in functional connectivity between those regions might be associated with the cognitive impairments observed in these DS mice.

First, we found that theta coherence between the hippocampus and mPFC was significantly greater in Dp1Tyb mice compared to WT littermates (Figure 4a), while no such differences were observed between Dp10Yey or Dp17Yey mice and their control groups (Figure 4b,c), or between WT cohorts (Supplementary Figure 2g). In addition, there were no differences in either low or high gamma coherence between hippocampus and mPFC in any mutant mouse group compared to their WT controls (Supplementary Figure 9). To confirm that the increased theta coherence observed in Dp1Tyb mice, compared to their WT littermates, did not simply arise due to the observed differences in movement statistics, we again removed any effect of average time spent immobile on theta coherence values across animals in both groups by linear regression, and then compared the residual values between groups (Supplementary Figure 6g,h). This confirmed that the increased theta coherence exhibited by Dp1Tyb mice was independent of differences in movement statistics.

**Figure 4:**
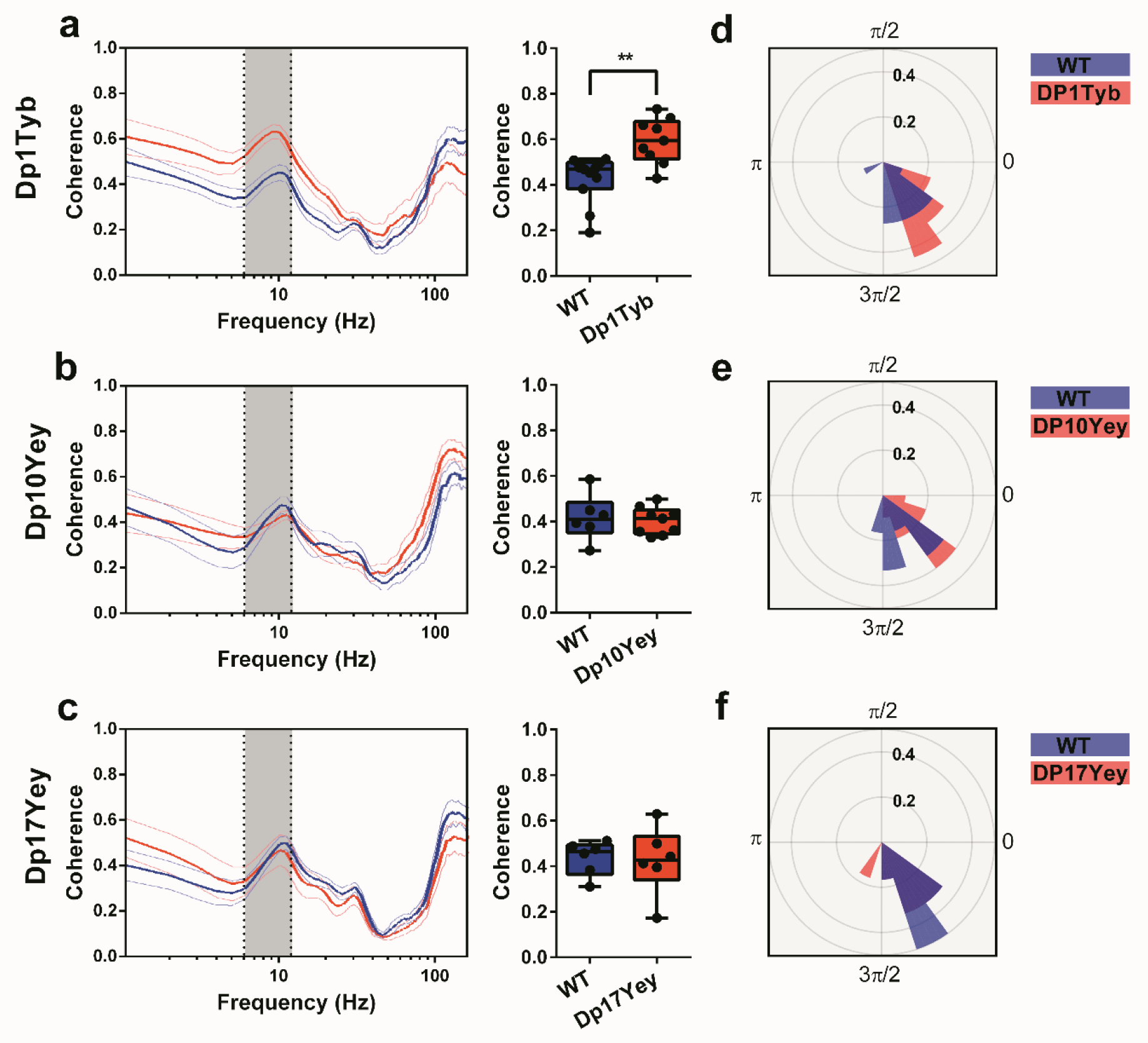
Hippocampal-medial prefrontal Phase Coupling during Spontaneous Alternation. **(a-c)** Coherence spectra and mean theta band coherence illustrating hippocampal-mPFC phase coupling during spontaneous alternation behaviour. Grey rectangles indicate the 6-12Hz theta band. **(a)** Theta band coherence is significantly greater in Dp1Tyb mice compared to WT (Mann-Whitney U=11.0, p<0.005), while there is no difference between either **(b)** Dp10Yey and WT or **(c)** Dp17Yey and WT animals. **(d-f)** Circular mean phase offset between mPFC and hippocampus for **(d)** Dp1Tyb and WT; **(e)** Dp10Yey and WT; and **(f)** Dp17yey and WT animals. The radial axis shows relative frequency, and the polar axis indicates the circular mean theta phase difference between mPFC and hippocampus. These results suggest that hippocampal theta oscillations lead those in mPFC by ∼1 radian (equivalent to ∼20ms at 9Hz) in all mutant and WT mice, without any differences between groups (Watson-Williams test, all p>0.07). Data are presented as box-whisker plots indicating the median, 25th and 75th percentiles, minimum and maximum values, with data for individual mice superimposed. Please refer to Supplementary Table 2 for full details of all statistical analyses.

To further characterise potential changes in functional connectivity across mouse lines, we extracted the theta phase lag between hippocampus and mPFC in order to assess the direction of communication between these regions (Figure 4d-f). In each group of animals, we found that hippocampal theta oscillations led those in mPFC by ∼1 radian, which is equivalent to ∼20ms for a 6-12Hz theta oscillation, without any difference between strains. Intriguingly, these results indicate that Dp1Tyb mice – which exhibit slowed planning and decision-making behaviour during the spatial alternation task – showed increased theta-band coherence between hippocampus and mPFC, without any differences in the direction of communication between those regions. This suggests that cognitive dysfunction arising from these electrophysiological differences is due to an increased influence of hippocampal inputs on medial prefrontal dynamics, rather than changes in the direction of information flow between regions.

### Behavioural and LFP Characteristics are preserved across the Lifespan in DS Mouse Models

Finally, we asked whether the behavioural and LFP abnormalities observed in Dp1Tyb and Dp10Yey mice persisted throughout life, or were specific to the adolescent period during which they were initially tested^59^. To this end, we repeated tests of alternation behaviour and recorded LFP data from the same animals at six and nine months of age, alongside age matched WT controls (see Supplementary Figure 1c for further details, Supplementary Table 3 for animal and trial numbers). Importantly, we found that the differences in both behaviour and neural dynamics described above remained stable throughout this long-term assessment period.

First, we found that trial latency was significantly greater in Dp1Tyb mice compared to their WT control group across all three time points (Figure 5a), and the observed reduction in both hippocampal and mPFC peak theta frequency also persisted with age (Figure 5b,c). Similarly, hippocampal theta-HG phase-amplitude coupling was significantly greater in Dp1Tyb mice compared to WT at all ages (Figure 5d); and theta coherence between hippocampus and mPFC remained significantly higher than WT across the lifespan (Figure 5e).

**Figure 5:**
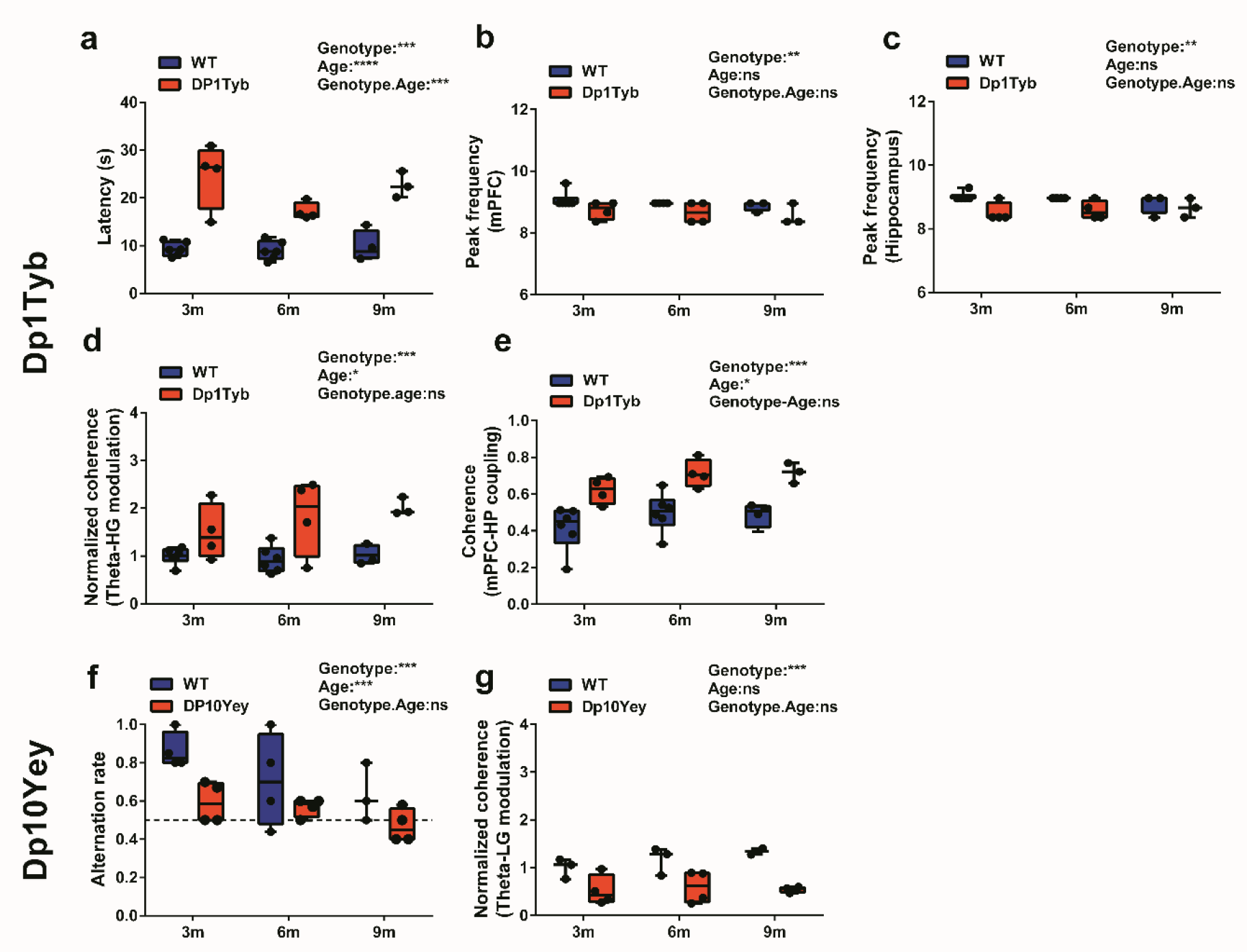
Behavioural and Electrophysiological Data across the Lifespan. Behavioural and LFP data at 3-4months (3m), 6-7 months (6m) and 9-10 months (9m) of age in: **(a-e)** Dp1Tyb; and **(f,g)** Dp10Yey mice. **(a)** Trial latency remains significantly greater in Dp1Tyb mice compared to WT throughout the lifespan (GLM, Type III tests χ2(1)=56.1, p<0.0001). Similarly, peak theta frequency in both **(b)** mPFC (GLM, Type III χ2(1)=6.84, p<0.01) and **(c)** hippocampus (GLM, Type III χ2(1)=8.93, p<0.01) is shifted to a significantly lower frequency. **(d)** Hippocampal theta-HG phase-amplitude coupling (GLM, Type III χ2(1)=14.2, p<0.0001) and **(e)** theta coherence between mPFC and hippocampus (GLM, Type III χ2(1)=29.6, p<0.0001) are also increased in Dp1Tyb mice at all three time points, compared to their WT control group. **(f)** Alternation rate remains significantly lower in Dp10Yey mice compared to WT throughout the lifespan (GLM, Type III χ2(1)=12.5, p<0.0001), and does not differ from chance level (black dashed line) at any age (Friedman’s test, χ2(5)=8.2, p>0.15), while the WT control group consistently perform above chance (Friedman’s χ2(5)=10.3, p<0.05). **(g)** Hippocampal theta-LG phase-amplitude coupling is also significantly lower in Dp10Tyb mice at all three time points (GLM, Type III χ2(1)=18.2, p<0.0001). Data are presented as box-whisker plots indicating the median, 25th and 75th percentiles, minimum and maximum values, with data for individual mice superimposed. Please refer to Supplementary Table 2 for full details of all statistical analyses.

Second, we found that the impaired alternation rate observed in young Dp10Yey mice persisted with age (Figure 5f). In contrast to WT mice, the alternation rate in Dp10Yey mice was not different from chance at any time point. Similarly, hippocampal theta-LG phase-amplitude coupling remained consistently lower in Dp10Yey mice relative to WT controls (Figure 5g). In sum, these results suggest that the observed differences in behaviour and neural dynamics between these DS mouse models and their WT control groups generally remained stable throughout adulthood, suggesting that aging neither alleviated nor worsened the phenotype in either strain.

## Discussion

The present study reveals distinct cognitive deficits and electrophysiological differences in three mouse models of DS which, combined, carry duplications covering all of the Hsa21-orthologous regions. By taking an unbiased approach, we aimed to discover if cognitive deficits resulting from triplication of genes/DNA elements in Hsa21 could be linked to individual regions with different sequence contents. As a measure of cognitive function, we used a canonical test of spatial memory – spontaneous alternation in a T-maze^60^. This behavioural test probes both decision-making and mnemonic function, based on the premise that mice have evolved an optimal strategy to explore their environment that relies on memorising previous trajectories and then using that information to plan future trajectories. Numerous cortical regions are implicated in successful performance of this task, most notably the hippocampus and mPFC^44, 61^.

Using this behavioural paradigm, we have shown that alternation deficits and hippocampal/mPFC neural dysfunction segregate with different regions of homology in the DS models. First, we found that alternation rate, a putative index of mnemonic function, was decreased in Dp10Yey mice. In contrast, trial latency, which provides an independent measure of cognitive processing that includes decision-making, planning, goal-directed behaviour, and attention^42, 43, 45^, was prolonged in Dp1Tyb mice. In addition, we have shown that Dp1Tyb mice have a lower peak frequency in the theta band in hippocampus and mPFC, an increase in phase-amplitude coupling between theta and high gamma in hippocampus, and a striking increase in theta phase coupling between mPFC and hippocampus – each of which is independent of the observed differences in movement statistics between Dp1Tyb animals and their WT littermates. Conversely, Dp10Yey mice exhibit decreased phase-amplitude coupling between theta and low gamma in the hippocampus; while Dp17Yey mice did not show any significant behavioral deficits in spatial alternation, or any alteration in the electrophysiology of hippocampus or mPFC. Crucially, the alterations in behavior and neural dynamics observed in our mutant mice are also unlikely to arise from differences in womb-environment, rearing or housing conditions, as we found no differences between WT littermate groups either behaviorally or physiologically.

Previous studies that have interrogated hippocampal function in similar mutant mouse populations^62–65^ have found no impairments in long-term spatial memory in Dp10Yey mice^62, 63^. In contrast, we observed decreased alternation rates in Dp10Yey mice suggestive of a spatial memory deficit^39^. These conflicting findings likely reflect subtle differences in the behavioral tasks employed, which emphasize complementary aspects of neural processing both within the hippocampus and among a wider network of functionally integrated brain regions, and should be the subject of further investigation^66–68^. The behavioral phenotype observed here was associated with a decrease in theta-gamma phase-amplitude coupling in the hippocampus. It has been well established that spatial memory relies on the periodic reactivation of encoded information by theta modulation of gamma oscillations in rodents^55, 69, 70^ and humans^71–75^, and so our finding of decreased gamma-theta coupling in Dp10Yey mice is consistent with their behavioral phenotype, and indicates specific abnormalities of hippocampal circuitry in this model. Our data may thus provide a functional basis for the memory problems evident in people with Down syndrome^7, 9, 10^. Dp10Yey mice were generated to carry an internal duplication spanning the 39 Hsa21 protein-coding orthologs mapping to Mmu10 and several of these genes, such as *ADAR2*, *S100B*, *CSTB*, *PRMT2*, and *TRPM2*, have been shown to play a role in brain development and function, such that aberrant gene dosage may be related to intellectual disability in DS^76, 77^.

An unexpected finding of this study was the delayed decision-making observed in Dp1Tyb mice with preserved memory function. Similar behavioral differences have also been found in humans with DS, who exhibit markedly slower reaction times^78–80^. This impairment has been attributed to deficits in executive function that involve information processing, attention, and inhibition^7, 9^, resulting in difficulty prioritizing, staying engaged with a task, and consistently responding in the same manner to certain situations^9, 81^. Importantly, we found that the increased trial latency observed in these animals was associated with a reduction in theta frequency across both hippocampus and mPFC. It is well established that theta frequency is correlated with running speed in rodents^82^, and so a potential explanation for both of these findings is that Dp1Tyb mice simply moved more slowly in general. However, although we found that Dp1Tyb mice spent more time immobile – presumably, reflecting their inability to retain focus on the task – they exhibited no differences in running speed compared to their WT littermates, and the observed reduction in theta frequency was still present when we restricted our analyses to movement periods only.

In addition, we found that the delayed decision-making in Dp1Tyb mice was associated with increased hippocampal-mPFC theta coherence. Communication between mPFC and hippocampus occurs through both direct projections and bidirectional pathways via intermediaries in the thalamus, perirhinal and lateral entorhinal cortices^83–85^. It is well accepted that coherence of neuronal activity across brain regions serves as a general mechanism for increasing effective communication during memory and attention tasks. Hippocampal-prefrontal theta-band synchrony facilitates hippocampal inputs to the mPFC and the integration of gamma-mediated cell assemblies in mPFC^26, 86^. In addition, theta-band synchrony has frequently been observed during spatial decision-making^35, 36^. Thus, our finding of increased hippocampal-mPFC theta coherence is consistent with the observed behavioral phenotype. Widespread increases in low frequency coherence between distributed brain networks, particularly including mPFC, are also observed in people with DS, are more evident in DS than in patients with other neurological disorders, and are inversely related to cognitive performance^15^.

*DYRK1A*, located on chromosome 21, is a major candidate protein-coding gene for several aspects of DS and encodes a kinase involved in neurodevelopment^46–51^. Overexpression of this gene in transgenic mice results in changes in inhibitory circuits in the mPFC^16, 52^ and may result in abnormal neural dynamics, particularly in the gamma band. Furthermore, *Dyrk1a* overexpression in mice induces learning and memory impairments detectable in the Morris water maze and Y-maze^52^. Here, we showed that reducing *Dyrk1a* to the normal two copies in Dp1Tyb mice failed to rescue the prolonged decision-making we observed in the spatial alternation task. Thus *Dyrk1a* overexpression is not required for this phenotype, leading us to conclude that another gene or genes, when present in three copies within the Dp1Tyb region, are involved in the abnormal decision-making behavior described here. This is an important result that may, in part, explain why most of the current competitive DYRK1A inhibitors fail to pass the pre-clinical stage with respect to improvement of cognitive impairments in DS^87^. Of the 148 protein coding genes within the region duplicated in Dp1Tyb mice, a handful are candidates for further exploration.

We note that there may also be critical effects from dosage sensitivity of non-protein coding elements on Hsa21 and our genetically unbiased approach will allow us to map to the DNA region, not to simply focus on the relatively limited set of protein coding elements for which we have functional information. Finding the genes (coding and non-coding) responsible for the cognitive and electrophysiological phenotypes observed in these mice has the translational potential to reveal important routes towards phenotype modifying therapies, for example, by antisense oligomers, but our results indicate that targeting a single gene is unlikely to be sufficient.

In summary, our study elucidates an important link between different regions of Hsa21, cognitive deficits and both local and long-range neural circuit dysfunction. Importantly, our results imply that specific cognitive deficits in Down syndrome may result from different underlying genetic, functional and regional abnormalities. This has important implications for understanding such cognitive deficits and indicates that therapies in Down syndrome will likely need to target multiple processes.

## Acknowledgements

EMCF and VLJT were supported by grants from the Wellcome Trust (grant numbers 080174, 098327 and 098328). VLJT was also supported by MRC programme U117527252 and by the Francis Crick Institute which receives its core funding from the UK Medical Research Council (FC001194), Cancer Research UK (FC001194) and the Wellcome Trust (FC001194). NB is supported by a Wellcome Trust Principal Research Fellowship (202805/Z/16/Z). We thank Eugene Yu and Mariona Arbones for mouse strains, and Nick Rawlins for comments on an earlier version of the manuscript. We also thank Dr Karen Cleverly and Ms Dorota Gibbins for colony management and genotyping and the Biological Research Facility of the Francis Crick Institute for animal husbandry.

## Author contributions

P.C. collected behavioural and electrophysiology data; P.C, D.B., N.S., analysed electrophysiology data; P.C., N.S., S.N. analysed histology data; F.K.W., V.L.J.T., E.M.C.F. provided mice for this study; S.S., M.G., T.C., N.B., M.C.W., E.M.C.F. helped direct the study; all authors contributed to the manuscript which was primarily drafted by P.C., D.B, M.C.W., E.M.C.F.

## STAR Methods

### Mouse Cohorts including Breeding and Ethics

We examined four mouse strains with the following alleles, previously described in^37, 38, 49^: C57BL/6J.129P2-Dp(16Lipi-Zbtb21)1TybEmcf/Nimr (hereafter referred to as Dp1Tyb); B6;129S7-Dp(10Prmt2-Pdxk)2Yey/J (hereafter referred to as Dp10Yey); B6.129S7 Dp(17Abcg1-Rrp1b)1Yey (hereafter referred to as Dp17Yey) and B6.129P2-*Dyrk1a*^tm1Mla^. Dp1Tyb, Dp10Yey, Dp17Yey animals were maintained within a facility at University College London, whereas mice for the Dp1Tyb × *Dyrk1a*^tm1Mla/+^ intercross were bred at the Francis Crick Institute, to generate Dp1Tyb*Dyrk1aKO mice in which both alleles were on the same chromosome following a genetic crossover. All strains were maintained in separate colonies as hemizygous mutants backcrossed for over ten generations to C57BL/6J, with age-matched WT littermates used as controls. All experiments were undertaken blind to genotype, which was decoded after experimental analysis and reconfirmed using an independent DNA sample isolated from post-mortem tail.

All experiments were performed in accordance with the United Kingdom Animal (Scientific Procedures) Act 1986. Reporting is based on the ARRIVE Guidelines for Reporting Animal Research developed by the National Centre for Replacement, Refinement and Reduction of Animals in Research, London, United Kingdom. Mice were housed in controlled conditions in accordance with guidance issued by the Medical Research Council in Responsibility in the Use of Animals for Medical Research (1993) and all experiments were carried out under License from the UK Home Office and with Local Ethical Review panel approval. Mice were housed in individually ventilated cages (IVC) of 2-5 age-matched animals under controlled environmental conditions (24–25°C; 50–60% humidity; 12 h light/dark cycle) with free access to food and water.

### Surgical Preparation and Transmitter Implantation for Long-term Recording

Mice were anaesthetised with 2.5-3 % isoflurane (Abbot, AbbVie Ltd., Maidenhead, UK) in 100% oxygen (flow rate of 1-1.5 litre/min) via gas anaesthesia mask (Model 906, David Kopf Instruments Tujunga, CA, USA) from a recently calibrated vaporizer (Harvard Apparatus, Cambridge, MA). Body temperature was maintained with a heat blanket during surgery. A transmitter (A3028A, Open Source Instruments, Brandeis, Boston, USA)^88^ was implanted subcutaneously with the depth recording electrodes (J-electrode, a teflon-insulated stainless steel electrode, Open Source Instruments, Brandeis, Boston, USA) positioned in mPFC (1.8 mm anterior, 0.4 mm lateral, 1.5 mm ventral) and dorsal hippocampus (1.85 mm posterior, 1.25 mm lateral, 1.45 mm ventral)^89, 90^. The reference electrode was implanted over the cerebellum posterior to lambda. The whole assembly was held in place with dental cement (Simplex Rapid, Acrylic Denture Polymer, UK). A subcutaneous injection of bupivacaine and metacam was provided for post-surgical pain management. At the end of surgery, enrofloxacin (5mg/kg, Baytril, Bayer health care) and pre-warm saline (0.5-1 ml) were administered subcutaneously. The animals were placed in a temperature controlled (25°C) recovery chamber until ambulatory and closely monitored at least 1-2 hours before returning to their home cage to allow recovery for at least 14 days after surgery.

The transmitter, which has no adverse effects^91^, was chronically implanted for longitudinal data recordings. During all recording sessions, continuous LFP recordings were recorded (bandpass filter: 0.2 Hz to 160 Hz, 512Hz sampling rate with 16 bit resolution) using LWDAQ Software (Open Source Instruments, Brandeis, Boston, USA). Animals were carefully monitored daily and were euthanized at the end of experiment with pentobarbital (25 mg/kg).

### Behavioural Testing: T-maze Spontaneous Alternation

Cognitive function in male mice from each strain and associated age-matched WT controls was assessed using the spontaneous alternation paradigm in an enclosed T-maze apparatus^39^. Animals were transferred to the testing room for 1-2 hours before each experiment to habituate to the environment and achieve an optimal state of arousal. Each mouse was then subjected to ∼10 trials per session, and sessions were completed at 3, 6, and 9 months of age (see Supplementary Tables 1 and 3 for average trial numbers in each group).

During each trial, the animal was first placed in the start chamber for 100s while reference phase LFP was recorded. Next, the guillotine door separating the start chamber from the central arm was raised and the mouse was allowed to run and choose a goal arm. After making a choice, the guillotine doors separating the central arm from each goal arm were slowly lowered, such that the animal was confined in the chosen goal arm which it could then explore for 30s. Next, the animal was transferred back to the start chamber, the guillotine door separating the central arm from the goal arms was raised and, after another 100s delay period in the start chamber, the guillotine door separating the start chamber from the central arm was raised again to allow the mouse a choice between the two open goal arms. Importantly, each trial included a free choice of goal arms on both the sample run and choice run (Figure 1a and Supplementary Figure 1a, b).

Trials were marked as successful if the mouse chose different goal arms on each run, and failures if the mouse chose the same goal arm on both runs. Alternation rate was defined as the total proportion of successful trials for each animal during each session. Trial latency was calculated as the time between the door isolating the start chamber being raised and the time at which the animals nose reached the decision point (i.e. exiting the central arm of the T-maze) immediately prior to the whole body completely entering the goal arm (indicative of a choice being made; see Figure 1b). This ensures that ‘vicarious trial and error’ behaviour, in which animals approach the decision point and look along either choice arm prior to making a decision, is excluded. Trial data was discarded if the latency on either run exceeded 120s.

Movement statistics were extracted from video data that covered the central arm, decision point, and initial stages of each goal arm, sampled at a rate of 25Hz, using the single mouse tracker plugin for Icy^92^. Running speed values were smoothed with a box car filter of 400ms width, and periods of immobility were defined as time frames when the animal’s movement speed was lower than 2cm/s.

### Histology

At the end of the experiment, the brain was removed and immediately immersed in 4% paraformaldehyde for >24 hours before being transferred to 30% sucrose post-fixation solution. Brain sections (40-μm thick thickness) were cut using a microtome (Leica SM2000R, Leica Microsystems ltd., United Kingdom) and stained with cresyl violet to allow histological location of the electrode track. This procedure allowed us to verify recording electrode locations, and LFP data were only included in the study if electrode tips were located in mPFC and dorsal hippocampus. In total, LFP data from just one animal was excluded because the recording site was outside the target region (Supplementary Figure 5).

### EEG Data Analysis

#### LFP Pre-processing

For our initial analyses, continuous LFP recordings from each region were segmented into 10s epochs that lasted from 5s before to 5s after animals reached the decision point on each run (plus 1s padding, subsequently discarded to account for potential edge effects). Each epoch was visually inspected for artefacts prior to further analysis using custom written Matlab (Mathworks, Natick MA) code (see Supplementary Tables 1 and 3 for trial numbers across strains). Trial latency and alternation rate data from trials excluded due to LFP artefacts were nonetheless included in behavioural analyses.

For subsequent analyses in which the relationship between movement statistics and EEG features were examined, continuous LFP recordings from each region were segmented to match the available movement data (plus 1s padding, subsequently discarded to account for potential edge effects). Any epochs that exhibited artefacts during visual inspection or for which video data (and therefore movement statistics) was either incomplete or unavailable were excluded from subsequent analysis.

#### Time-frequency Analysis

After de-trending and de-meaning the LFP signal from each trial, time-frequency decomposition was performed using a five cycle complex Morlet wavelet transform, with 1s of data from the beginning and end of each epoch subsequently discarded to avoid edge effects. Time-frequency representations were then averaged across this time window to provide a power spectrum for each epoch, and each power spectrum was then normalised by its integral to facilitate comparisons between animals. Finally, these normalised power values were averaged across the 6-12Hz theta band to provide an index of theta power in each epoch for statistical comparison.

In addition, to characterise the relationship between theta power and movement, we zero-phase filtered each LFP signal in the 6-12Hz theta band using a 400^th^ order finite impulse response (FIR) filter, discarded 1s of data from the beginning and end of the signal to avoid edge effects, extracted the analytic signal using the Hilbert transform, and then computed dynamic power and frequency. Running speed data was up-sampled to match the power and frequency time series, allowing us to compute average theta power during movement periods only and to estimate the intercept and slope of the running speed v theta frequency relationship in each animal using linear regression.

#### Phase-amplitude Coupling Analysis

To assay phase–amplitude coupling in the hippocampal LFP signal, we first computed cross-frequency coherence across a range of phase and amplitude frequencies following^54^. To do so, we extracted the amplitude at each time point across a frequency range of 20-160Hz from the Morlet wavelet transform described above, and then computed coherence between the original LFP signal and each of these amplitude time series across a phase frequency range of 2-40Hz using a window size of 1s and an overlap between subsequent windows of 0.5s. These coherence spectra subsequently index phase-amplitude coupling (PAC) between low frequency phase and high frequency amplitude, and can be aggregated across amplitude frequencies to generate the cross-frequency coherence images shown in Figure 3 and Supplementary Figure 7.

Visual inspection of cross-frequency coherence images averaged across all animal groups (shown in Supplementary Figure 7) revealed that 6-12Hz theta phase modulated the amplitude of higher frequency oscillations in two distinct bands, 60-120Hz (hereafter referred to as ‘low gamma’, LG) and 140-160Hz (hereafter referred to as ‘high gamma’, HG). We subsequently characterised the magnitude of theta-LG and theta-HG PAC in each epoch by zero-phase filtering the LFP signal separately in the 6-12Hz theta, 60-120Hz LG and 140-160Hz HG bands using a 400^th^ order FIR filter, extracting the analytic signal in each band using the Hilbert transform, and then computing the mean amplitude of the higher frequency oscillations in each of 30 evenly distributed theta phase bins. The resulting vector length of each mean amplitude distribution, computed using the circular statistics toolbox for Matlab^93^, provides an index of theta-LG and theta-HG PAC in each epoch for statistical comparison.

#### Phase Coupling Analysis

To compute an index of theta phase coherence between LFP recordings from the hippocampus and mPFC in each epoch, we first generated coherence spectra for each epoch using a window size of 1s and an overlap between subsequent windows of 0.5s and then averaged coherence values across the 6-12Hz theta range. In addition, to estimate the theta phase lag between concurrent oscillations in these regions, we zero-phase filtered each LFP signal in the 6-12Hz theta band using a 400^th^ order FIR filter, discarded 1s of data from the beginning and end of the signal to avoid edge effects, extracted the analytic signal using the Hilbert transform, and then computed the circular mean theta phase difference between regions across all time points within each epoch. This provides an indication of the time lag between those signals in the 6-12Hz theta band (computed by dividing the phase difference by the angular frequency at the centre of the theta band, i.e. 18π rad/s).

#### Correcting for Differences in Movement Statistics

Where significant differences in movement statistics between groups existed, we attempted to eliminate any potential confound on concomitant differences in theta coherence and theta-gamma PAC by linear regression. Specifically, we extracted the residual coherence or PAC values after regressing the amount of time spent immobile against those parameters across all animals (mutant and WT), and then assessed the difference in residual values between groups.

#### Statistical Analysis

Detailed statistical analysis was performed using SPSS 24 (Statistical Product and Service Solutions, IBM). All data are presented as mean ± SEM. Comparisons of means were performed using two-tailed Student’s t test and one way ANOVA with Tukey post hoc test if the data were normally distributed; Wilcoxon Signed test, Friedman’s test, or Mann-Whitney U-test if the data were not normally distributed (with the Shapiro-Wilk test and Kolmogorov-Smirnov test with Lillefors correction used to assess normality of the data distributions). Generalized linear model (GLM) Type III tests followed by Bonferroni post hoc tests were used for analysis of repeated measures longitudinal data. For circular (i.e. phase lag) data, the Watson-Williams test was used to assess differences between groups^93^. Differences were considered statistically significant at p<0.05. For full details of all statistical analyses, please refer to Supplementary Table 2.

## Supplementary Information

**Supplementary Figure 1:**
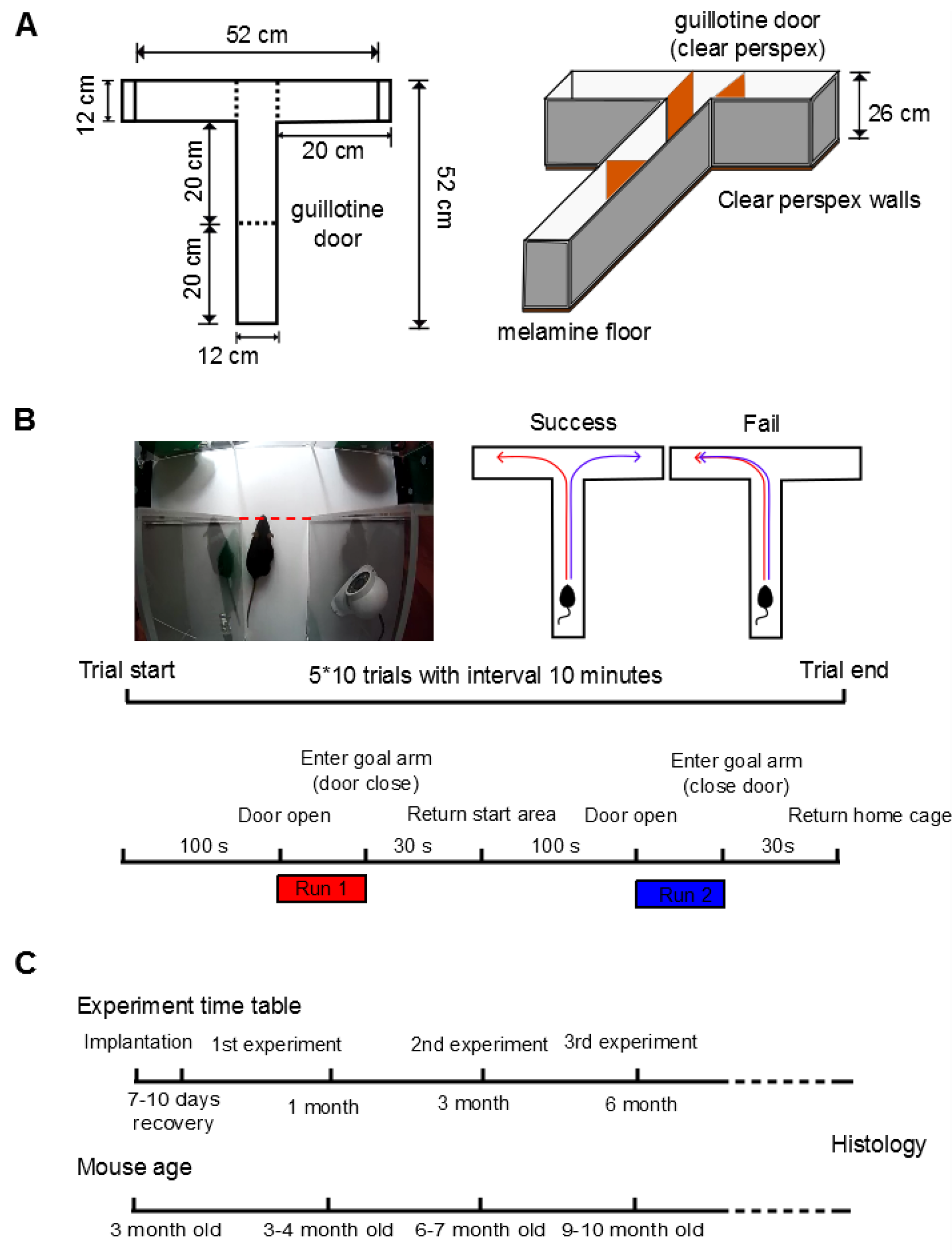
Further Details of the Behavioural Protocol. Experimental protocol for the T-maze spontaneous alternation task, showing: **(a)** a schematic of the T-maze; **(b)** the trial protocol for probing spontaneous alternation behaviour maze; and **(c)** a time line for the longitudinal study.

**Supplementary Figure 2:**
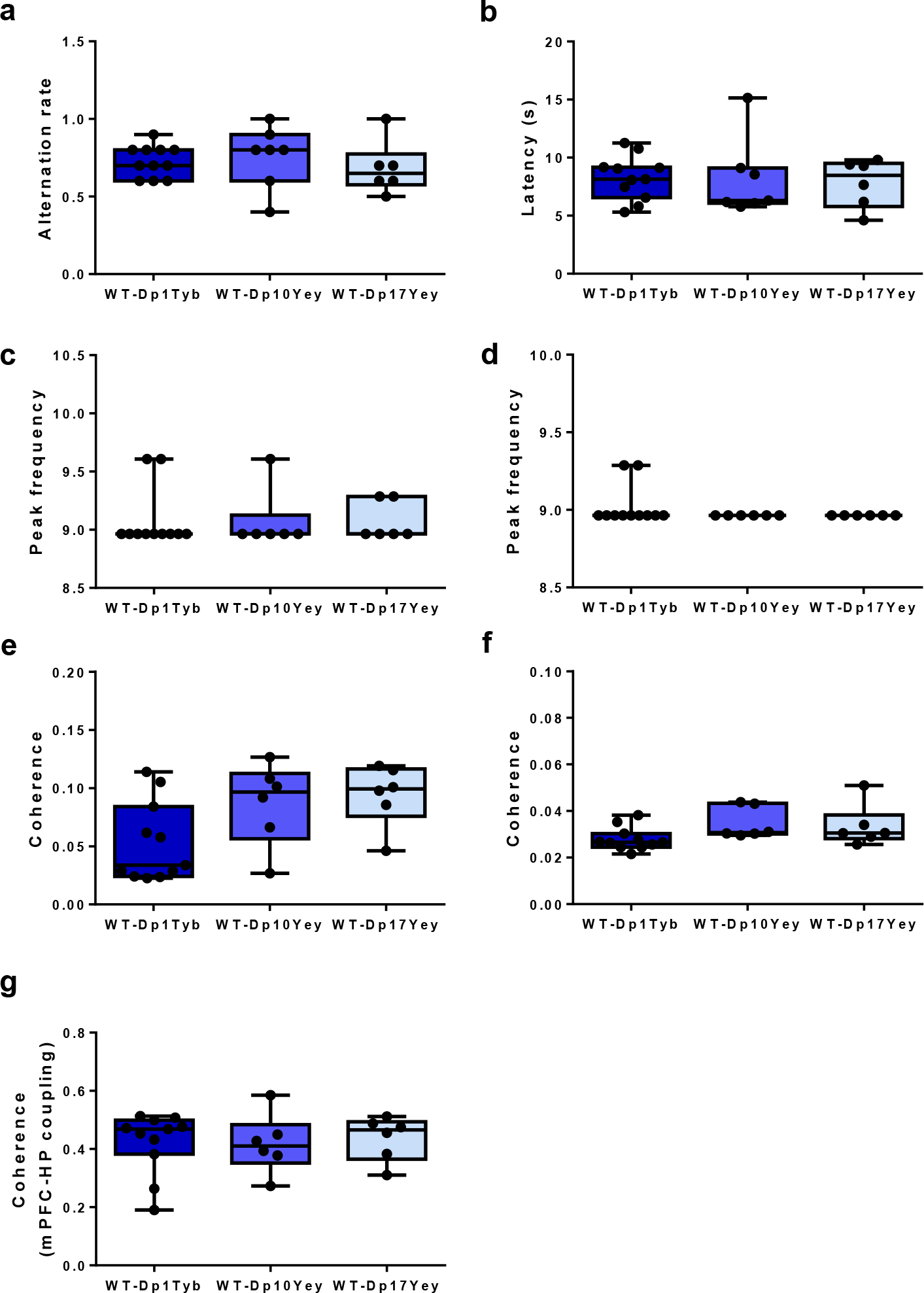
Comparison of behavioural and physiological data across WT groups. Comparison of **(a)** spatial alternation rate; **(b)** trial latency; peak theta frequency in **(c)** hippocampus and **(d)** medial prefrontal cortex (mPFC); **(e)** theta-low gamma and **(f)** theta-high gamma phase-amplitude coupling in the hippocampus; and **(g)** theta coherence between hippocampus and mPFC across WT cohorts. Data are presented as box-whisker plots indicating the median, 25th and 75th percentiles, minimum and maximum values, with data for individual mice superimposed. There are no significant differences between WT groups in any panel. Please refer to Supplementary Table 2 for full details of all statistical analyses.

**Supplementary Figure 3:**
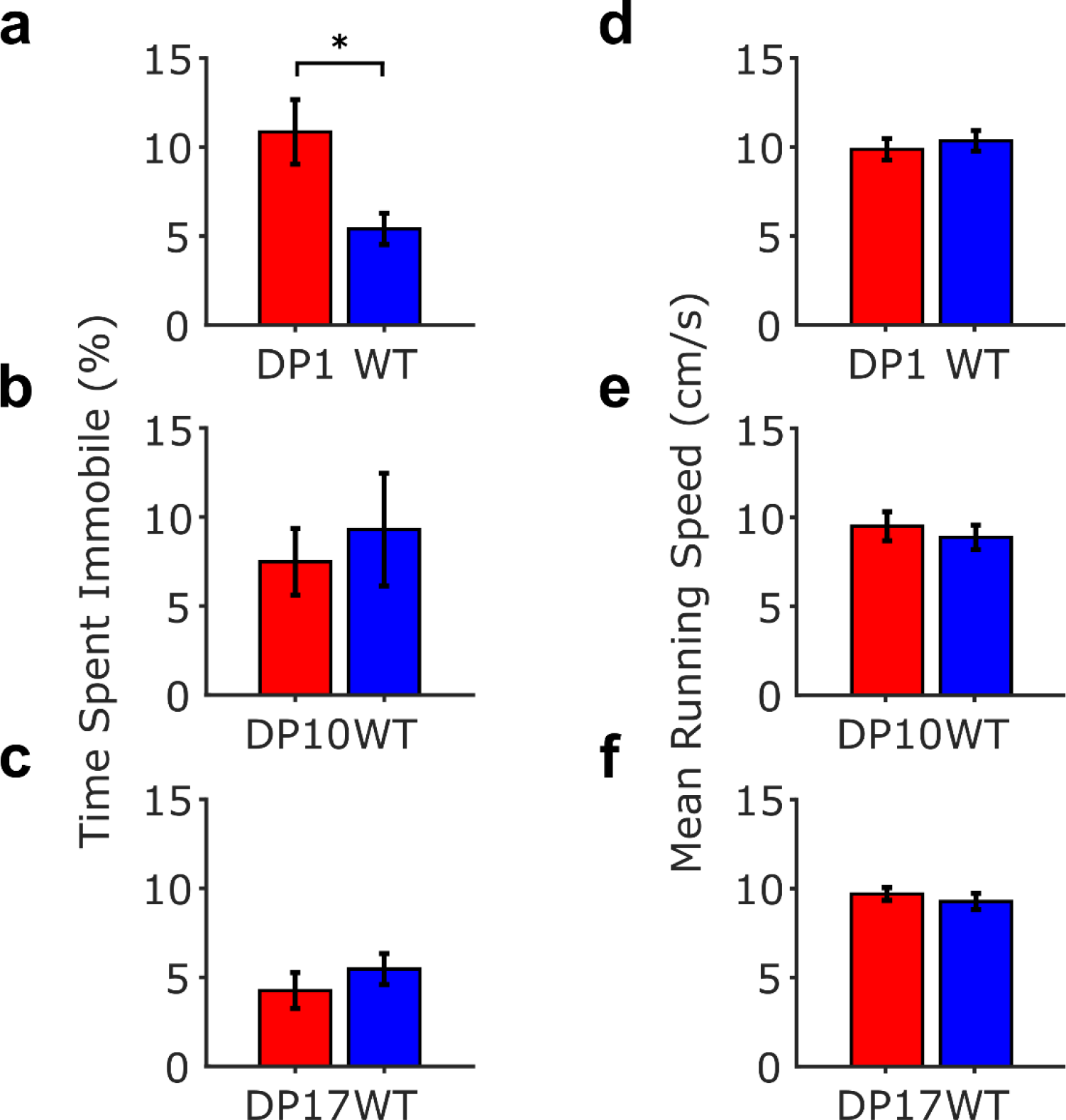
Comparison of movement statistics across groups (a-c) Relative period of each trial spent immobile (i.e. running speed <2cm/s); and **(d-f)** the average running speed during movement (i.e. running speed ≥2cm/s) across all mutant mouse and WT control groups. These data illustrate that the increased trial latency observed in Dp1Tyb mice results from significantly more time spent immobile (t(13)=2.46, p<0.05) without any difference in mean running speed during movement (t(13)=-0.49, p=0.63). No differences in time spent immobile or mean running speed during movement were observed in any other group compared to their WT cohort (all p>0.38), or between any of the WT groups (both p>0.24). Please refer to Supplementary Table 2 for full details of all statistical analyses.

**Supplementary Figure 4:**
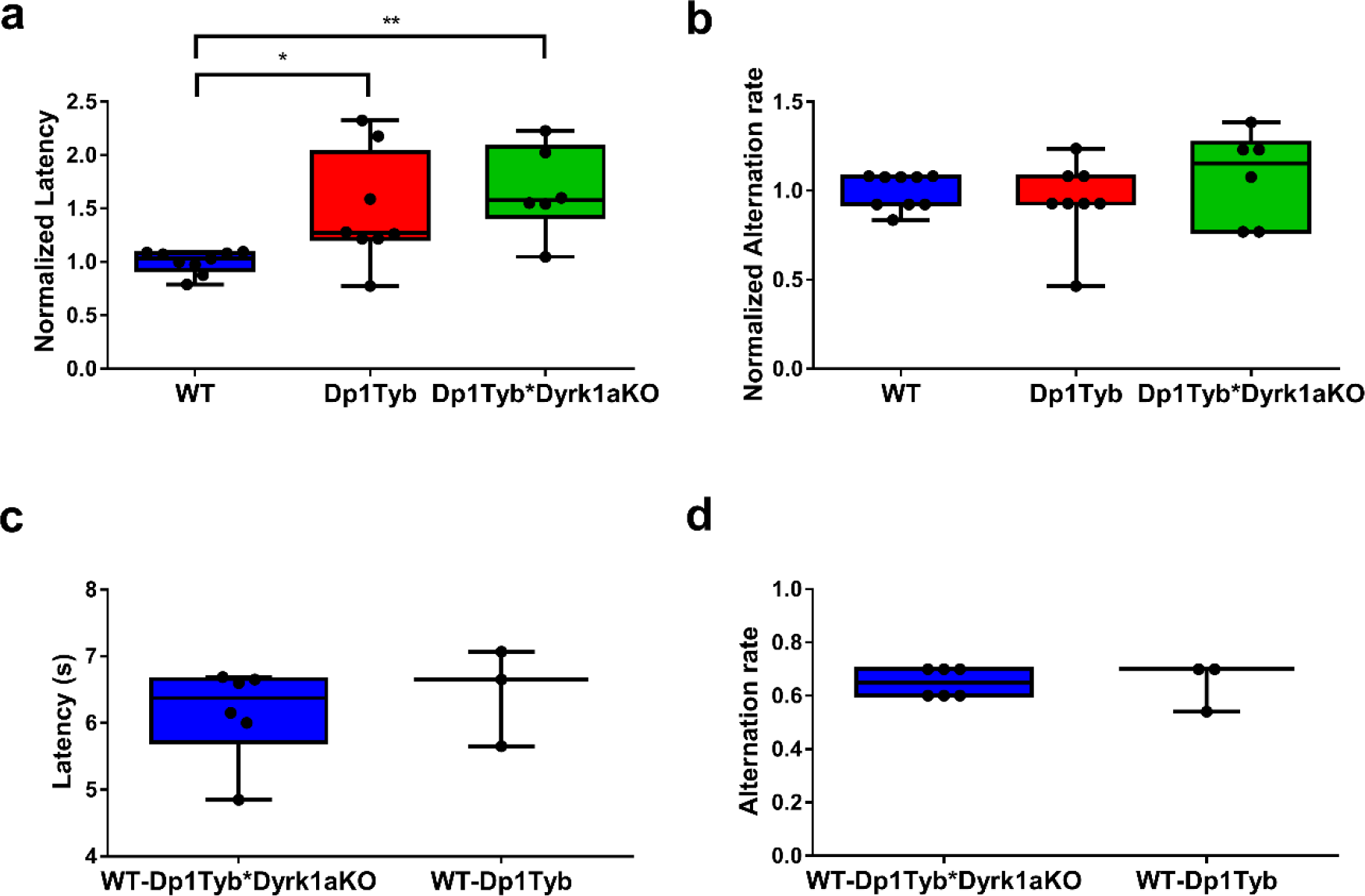
Behavioural performance in Dp1Tyb and Dp1Tyb*Dyrk1aKO mice. **(a)** Alternation rate and **(b)** trial latency averaged over 10 trials for Dp1Tyb and Dp1Tyb*Dyrk1aKO mice compared to their pooled wild-type (WT) control group. All data are normalized to the WT mean. Trial latency differs significantly between groups (One way ANOVA, F(2,20)=6.21, p<0.0001). Post-hoc Tukey HSD indicates that trial latency is significant longer in Dp1Tyb and Dp1Tyb*Dyrk1aKO mice, compared to pooled WT (p<0.05 and p<0.01, respectively). **(c)** Comparison of trial latency and **(d)** alternation rate between WT cohorts, illustrating no significant differences in either case (Kruskal-Wallis test, both p>0.58). Data are presented as box-whisker plots indicating the median, 25th and 75th percentiles, minimum and maximum values, with data for individual mice superimposed. Please refer to Supplementary Table 2 for full details of all statistical analyses.

**Supplementary Figure 5:**
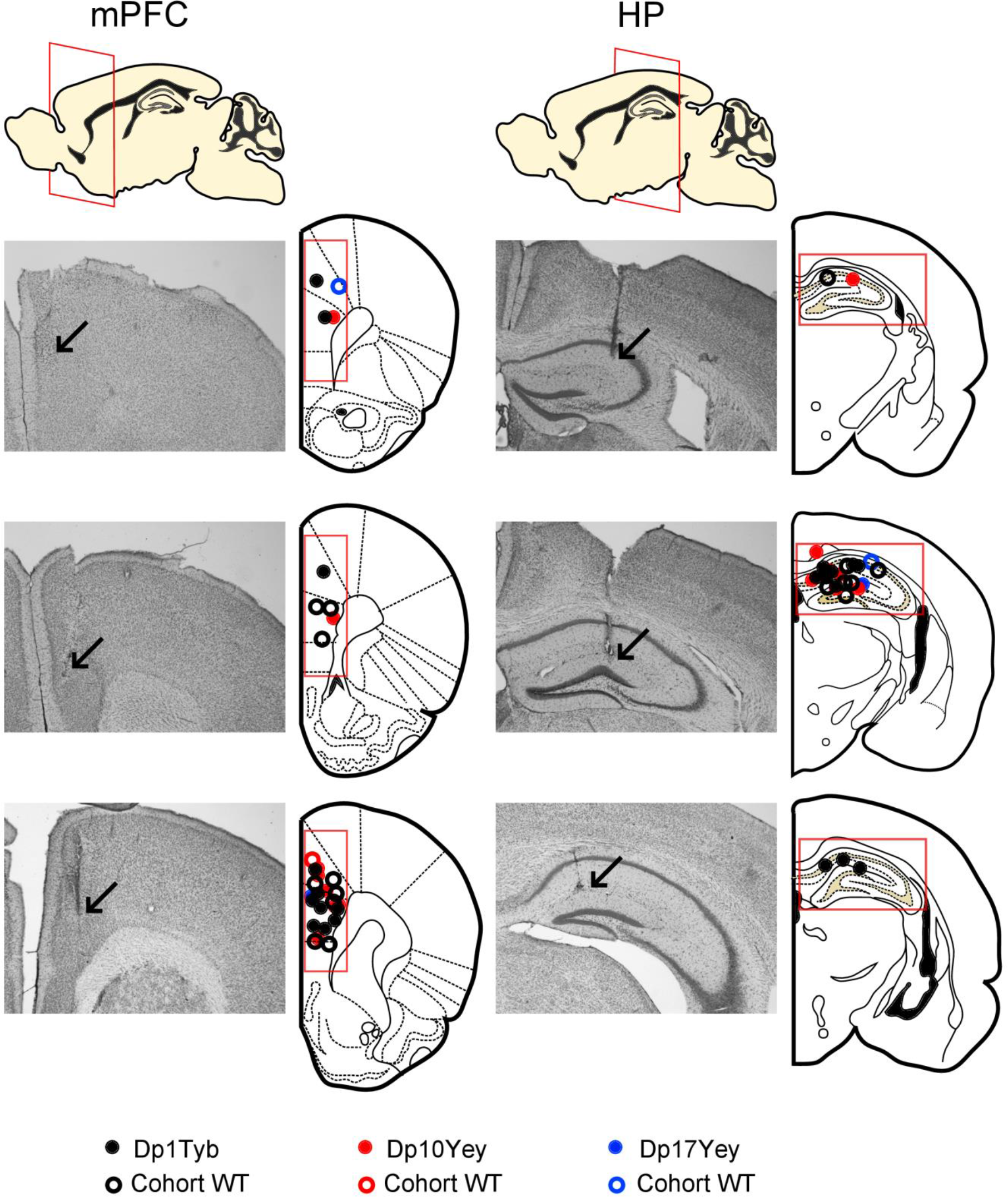
Histology. Coronal sections of Nissl stained brains showing typical locations of recording electrode tips in medial prefrontal cortex and dorsal hippocampus. Arrows indicate the tip of each recording electrode. LFP data were only included in the study if electrode tips were located in mPFC (indicated by coloured rectangle) and dorsal hippocampus (HP, also indicated by coloured rectangle).

**Supplementary Figure 6:**
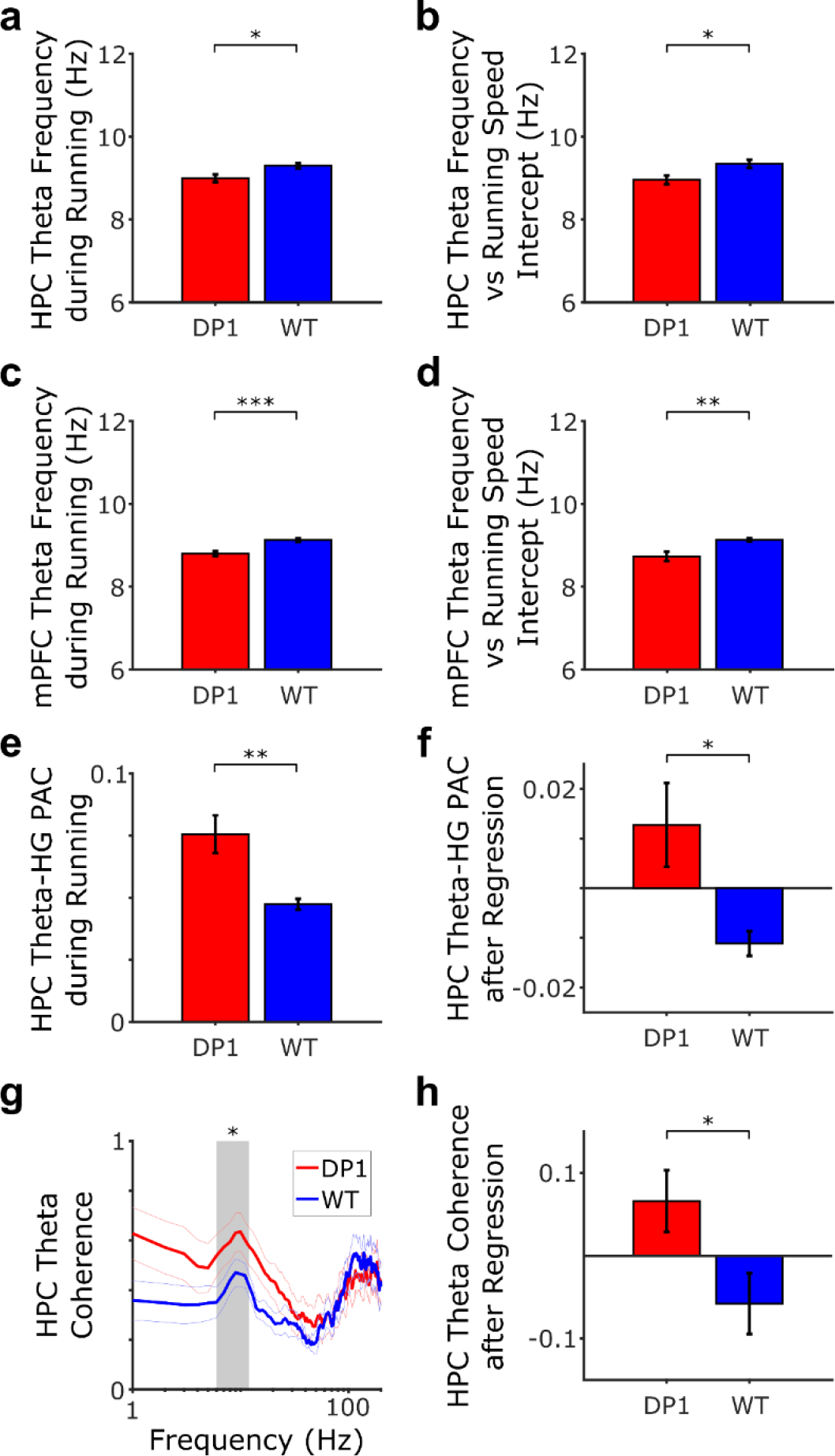
Details of Dp1Tyb analyses controlling for movement statistics. **(a,c)** Average theta frequency and **(b,d)** intercept of the running speed v theta frequency relationship during movement for Dp1Tyb and WT groups in **(a,b)** hippocampus (HPC) and **(c,d)** medial prefrontal cortex (mPFC). Theta frequency during movement is significantly lower in Dp1 animals in both HPC (t(13)=-2.80, p<0.05) and mPFC (t(13)=-4.52, p<0.001), due to a reduction in the intercept (HPC: t(13)=- 2.68, p<0.05; mPFC: t(13)=-3.45, p<0.01) but not the slope (both p>0.24, data not shown) of the running speed v theta frequency relationship. **(e)** Average theta-HG PAC in HPC is significantly higher in Dp1 animals when analyses are restricted to movement periods only (t(13)=3.12, p<0.01). **(f)** Moreover, the influence of average time immobile on theta-HG PAC is removed by linear regression across animals, the difference between groups is still significant (t(13)=2.88, p<0.05). **(g)** Average theta coherence between HPC and mPFC is significantly higher in Dp1 animals when analyses are restricted to movement periods only (t(13)=2.44, p<0.05). **(h)** Moreover, if the influence of average time immobile on theta coherence is removed by linear regression across animals, the difference between groups is still significant (t(13)=2.36, p<0.05). Please refer to Supplementary Table 2 for full details of all statistical analyses.

**Supplementary Figure 7:**
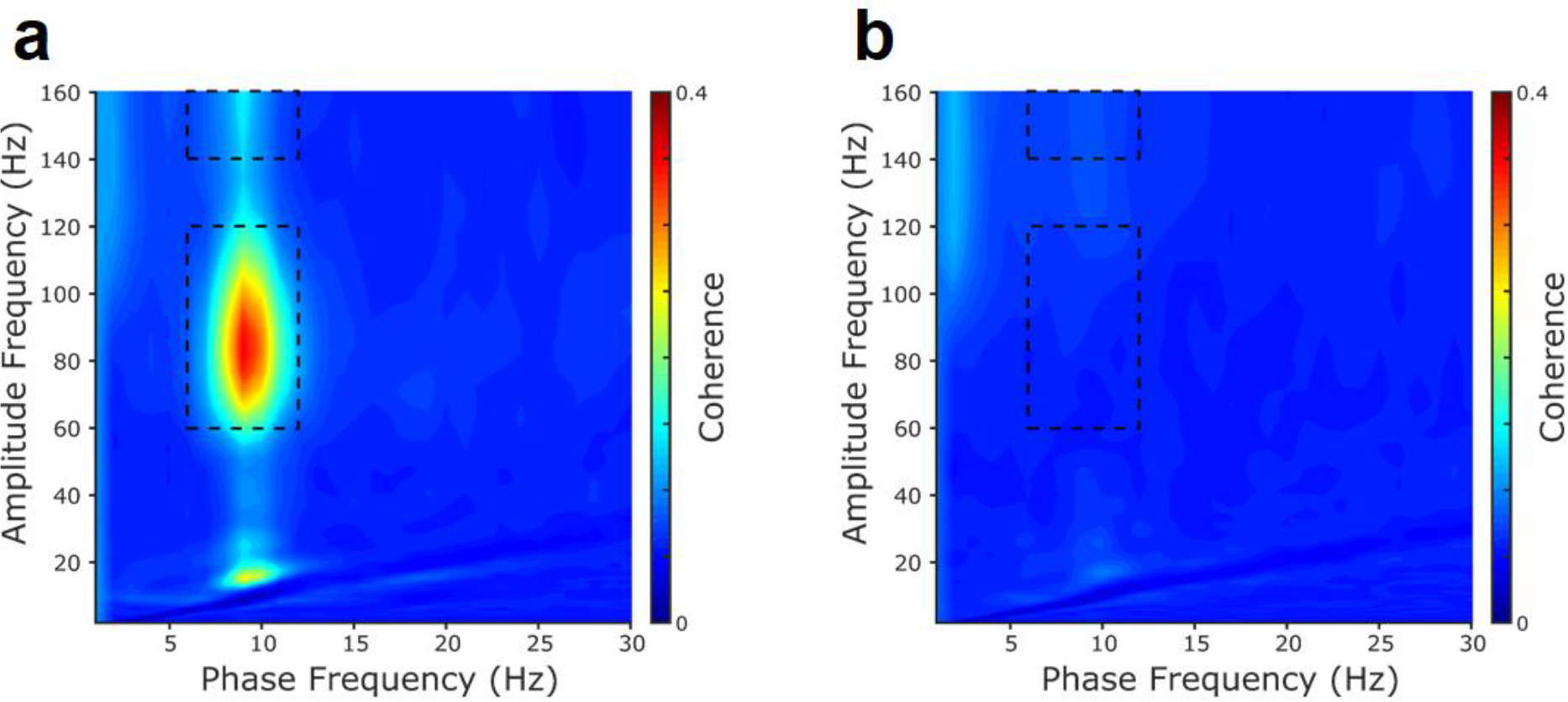
Average phase-amplitude comodulograms across all animals. Phase-amplitude comodulograms, averaged across all DS model and WT control groups at 3 months of age for **(a)** hippocampus and **(b)** mPFC. Visual inspection reveals strong peaks between 6-12Hz theta phase and both 60-120Hz ‘low gamma’ (LG) and 140-160Hz ‘high gamma’ (HG) rhythms in the hippocampus, with no strong phase-amplitude coupling apparent in the mPFC (see Figure 3 for details of comparisons between groups).

**Supplementary Figure 8:**
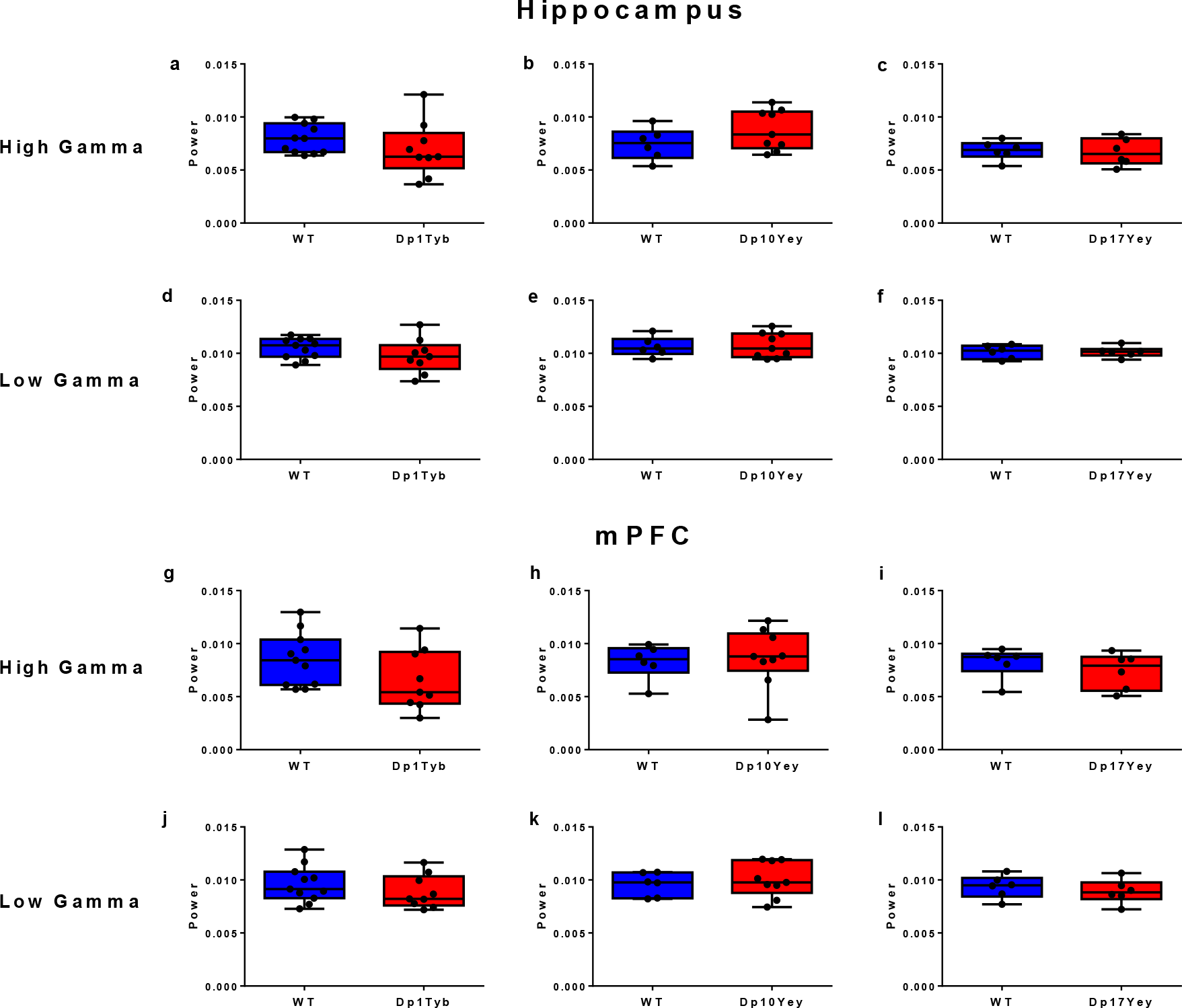
Comparison of Gamma Power between Groups. Average power in the low (60-120Hz) and high (120-140Hz) gamma bands in **(a-f)** hippocampus and **(g-l)** mPFC for **(a,d,g,j)** Dp1Tyb and WT; **(b,e,h,k)** Dp10Yey and WT; **(c,f,i,l)** Dp17Yey and WT animals during spontaneous alternation on the T-maze, which show no significant differences between mutant mice and WT groups in any instance. Data are presented as box-whisker plots indicating the median, 25th and 75th percentiles, minimum and maximum values, with data for individual mice superimposed. Please refer to Supplementary Table 2 for full details of all statistical analyses.

**Supplementary Figure 9:**
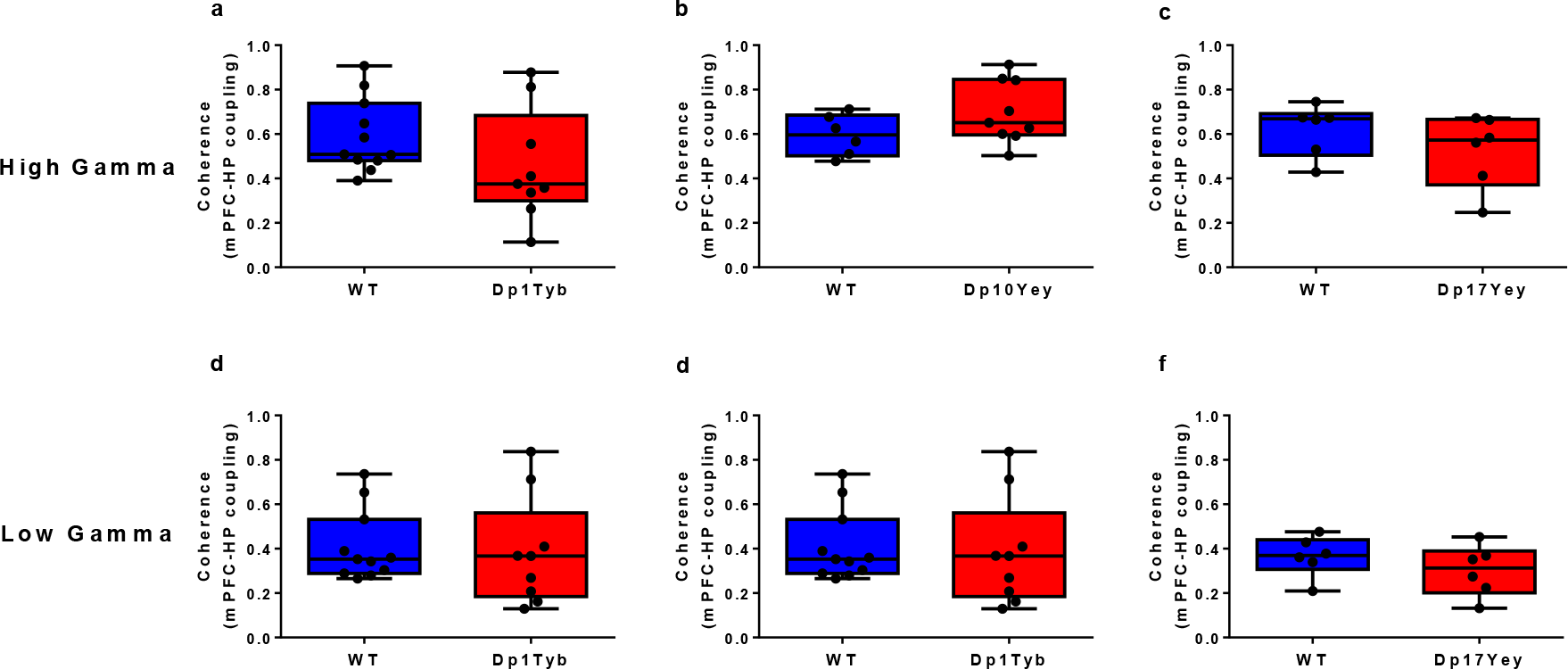
Comparison of Gamma Coherence between Groups. Hippocampal-medial prefrontal coherence in the **(a-c)** high- and **(d-f)** low-gamma frequency bands for each mutant mouse and WT control group, which show no significant differences in any instance. Data are presented as box-whisker plots indicating the median, 25th and 75th percentiles, minimum and maximum values, with data for individual mice superimposed. Please refer to Supplementary Table 2 for full details of all statistical analyses.

**Supplementary Table 1:**
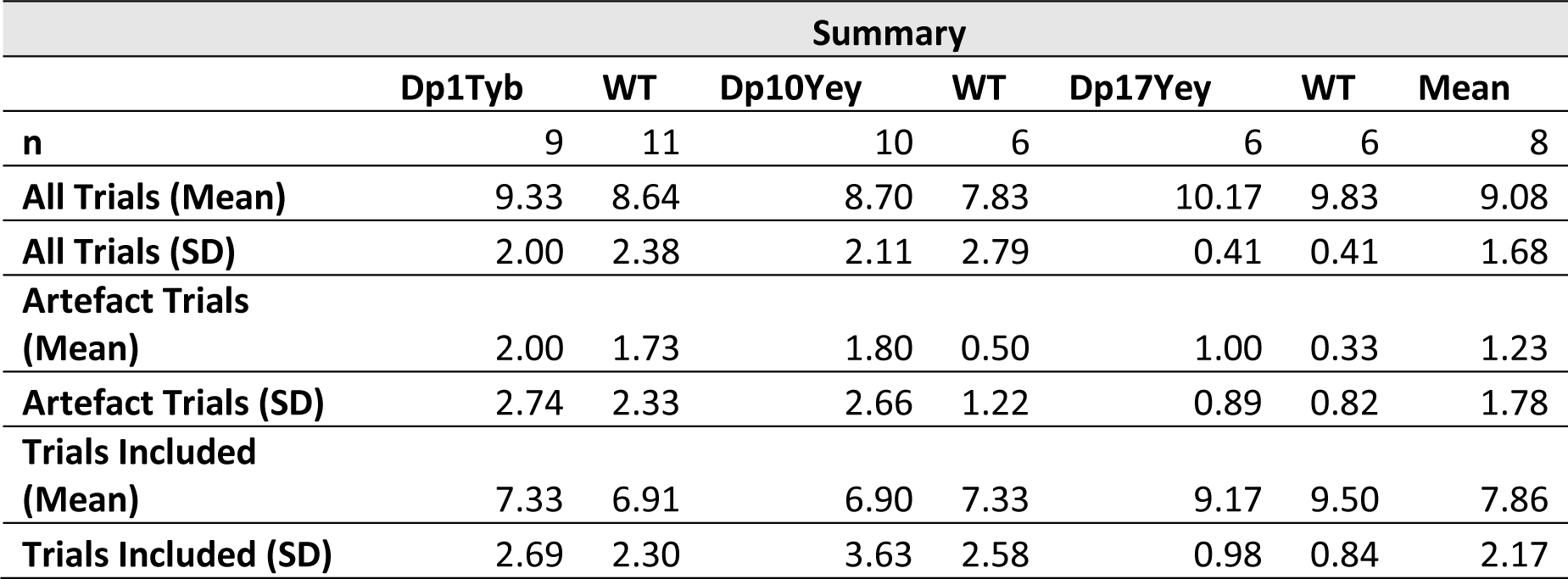
Summary of Trial and Animal Numbers at 3 months of Age

**Supplementary Table 2:**
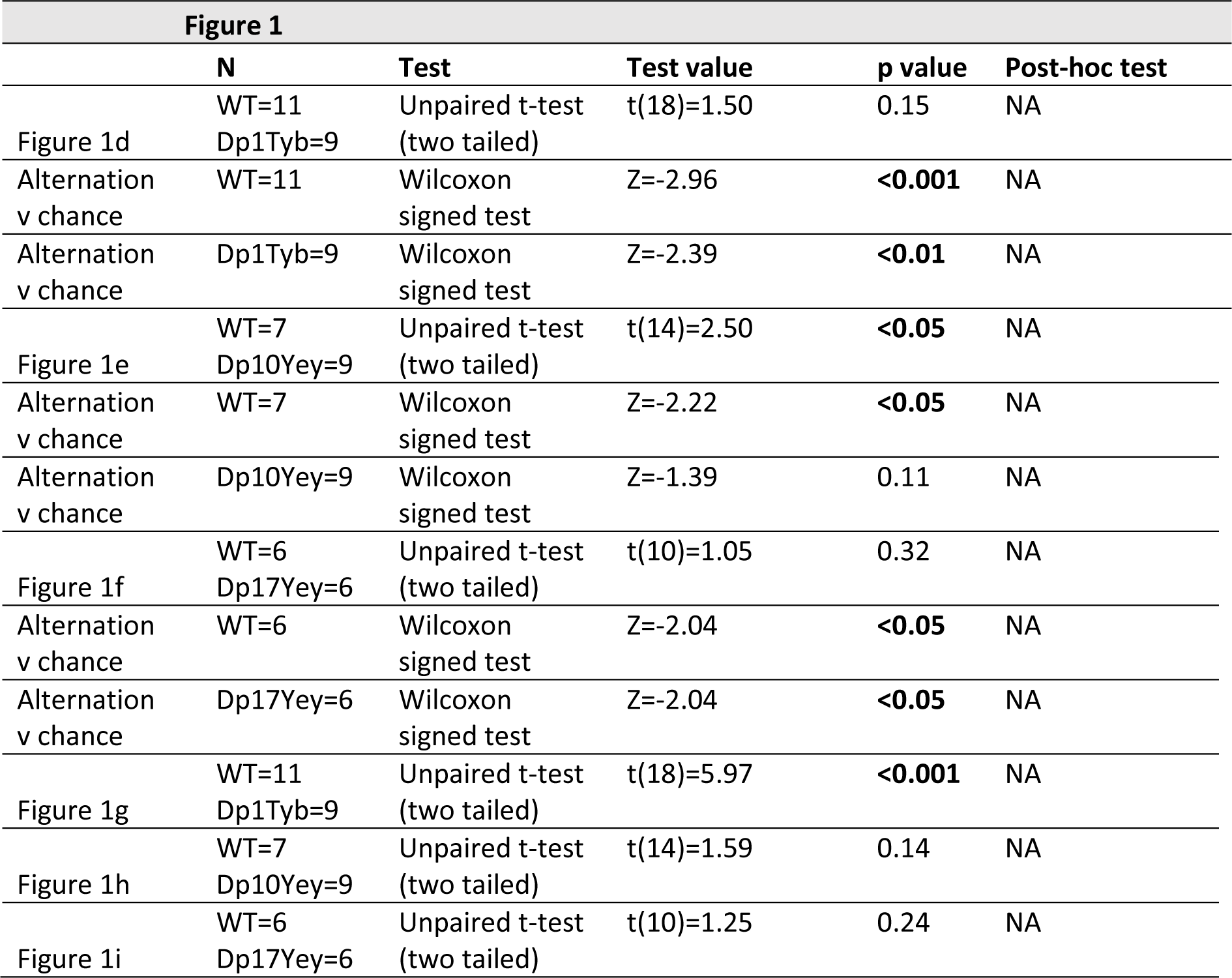
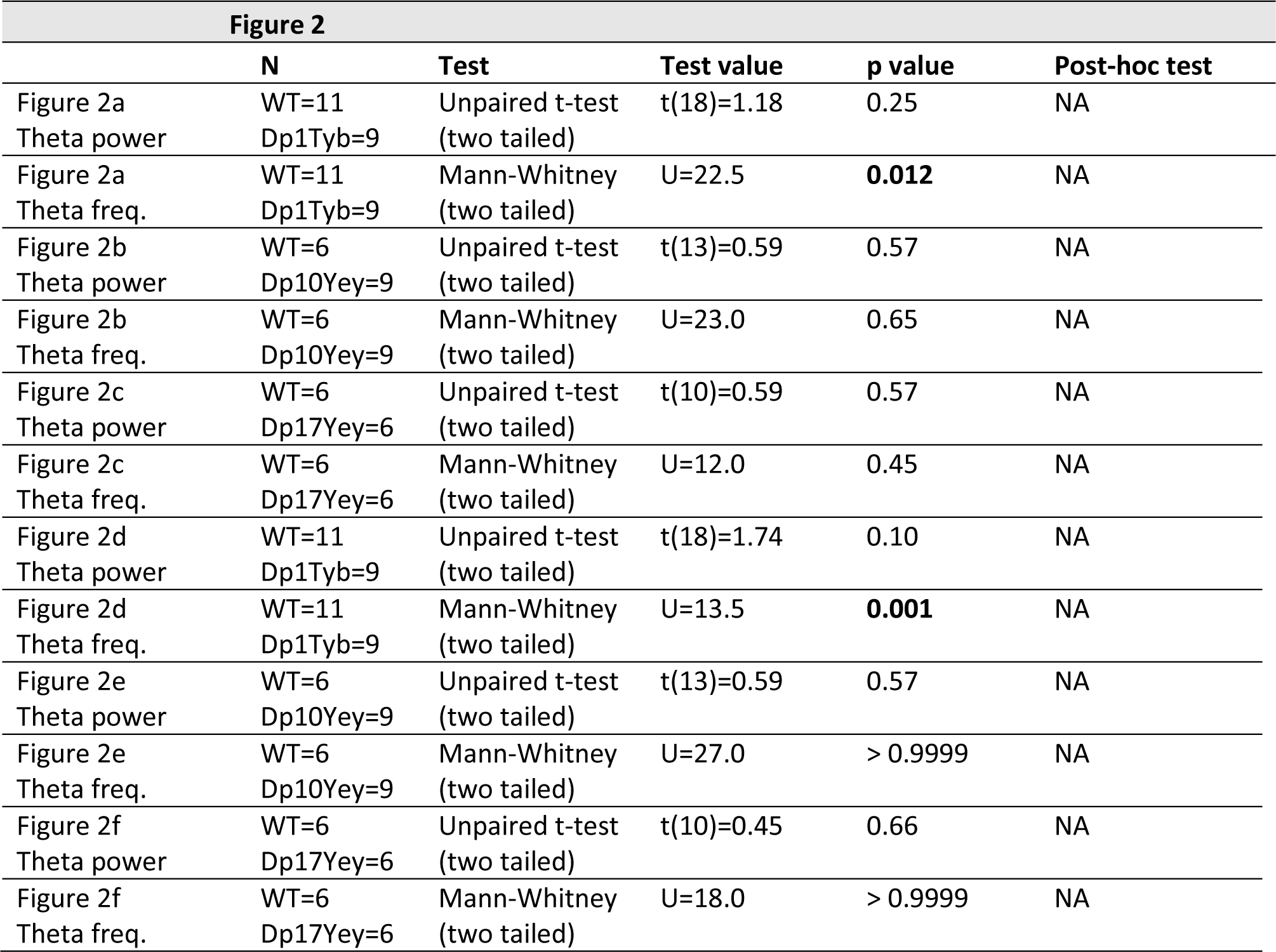
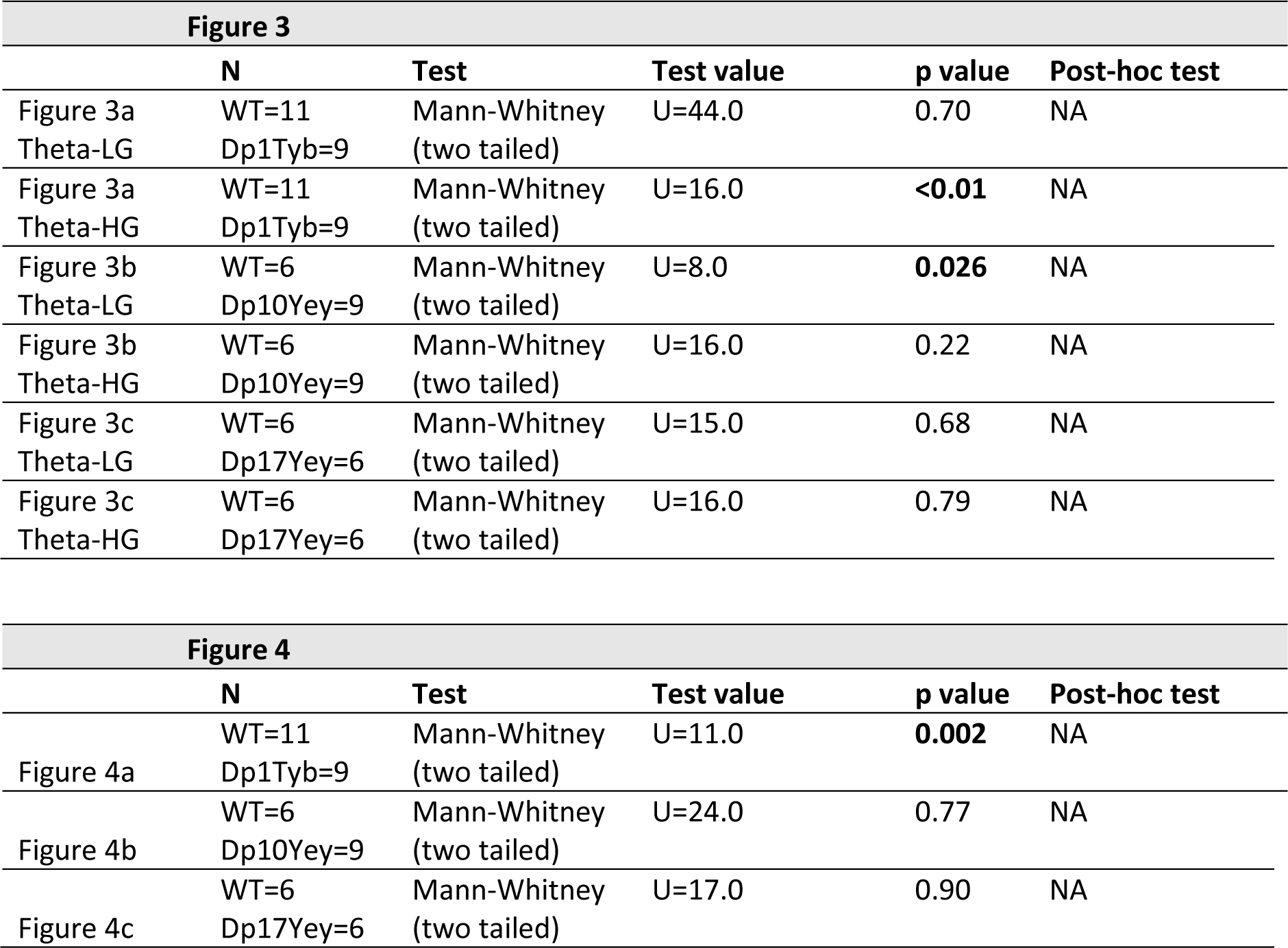
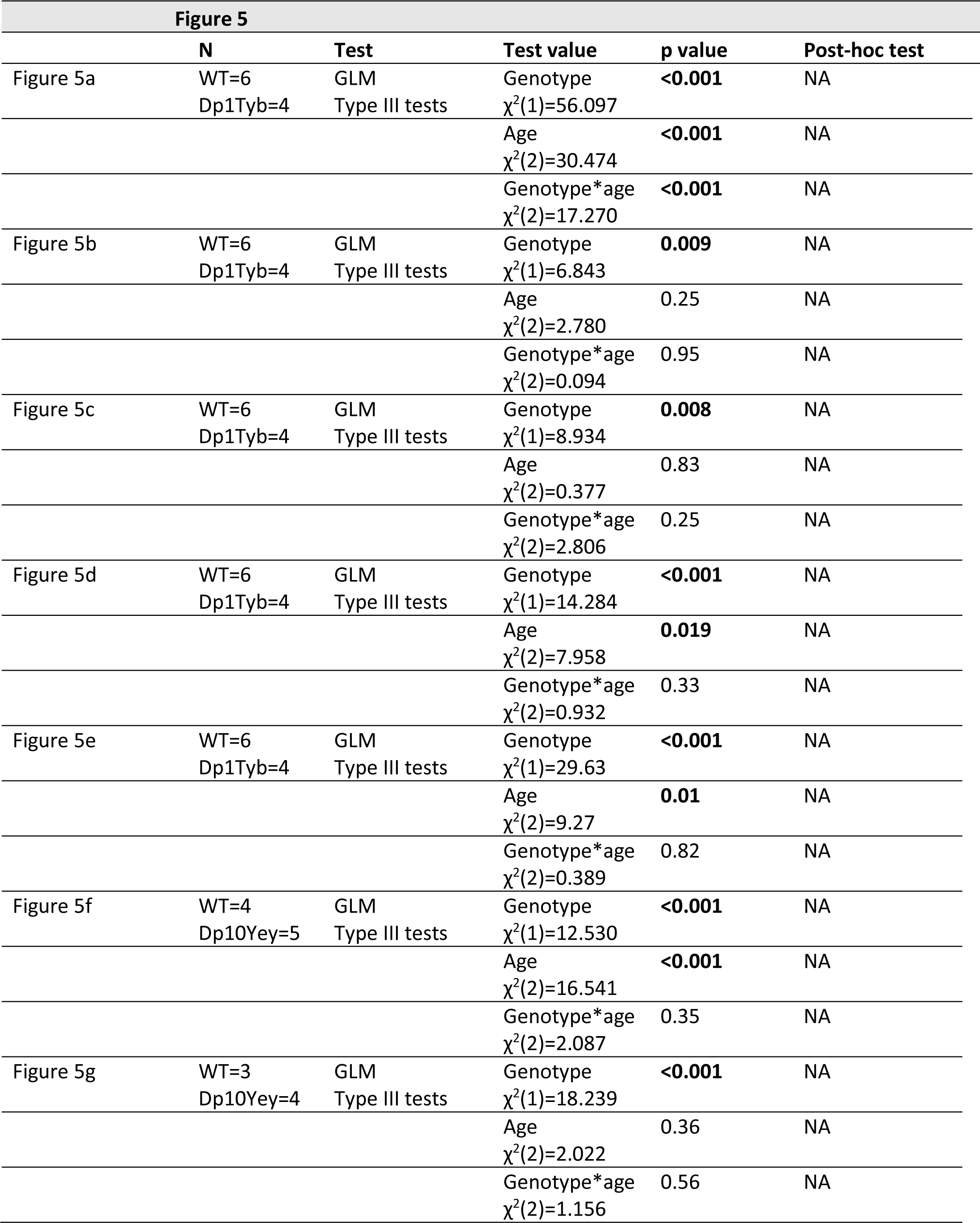
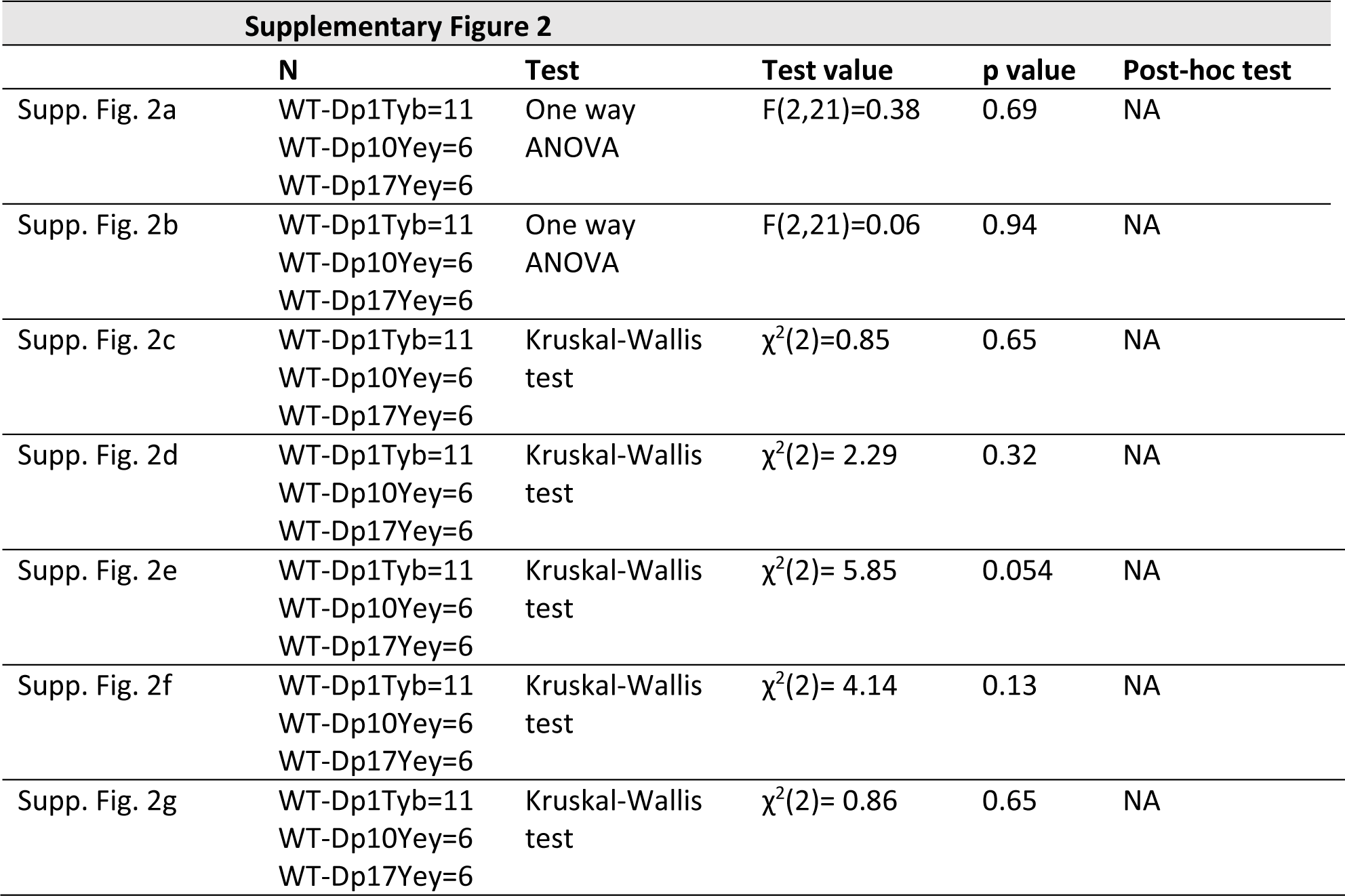
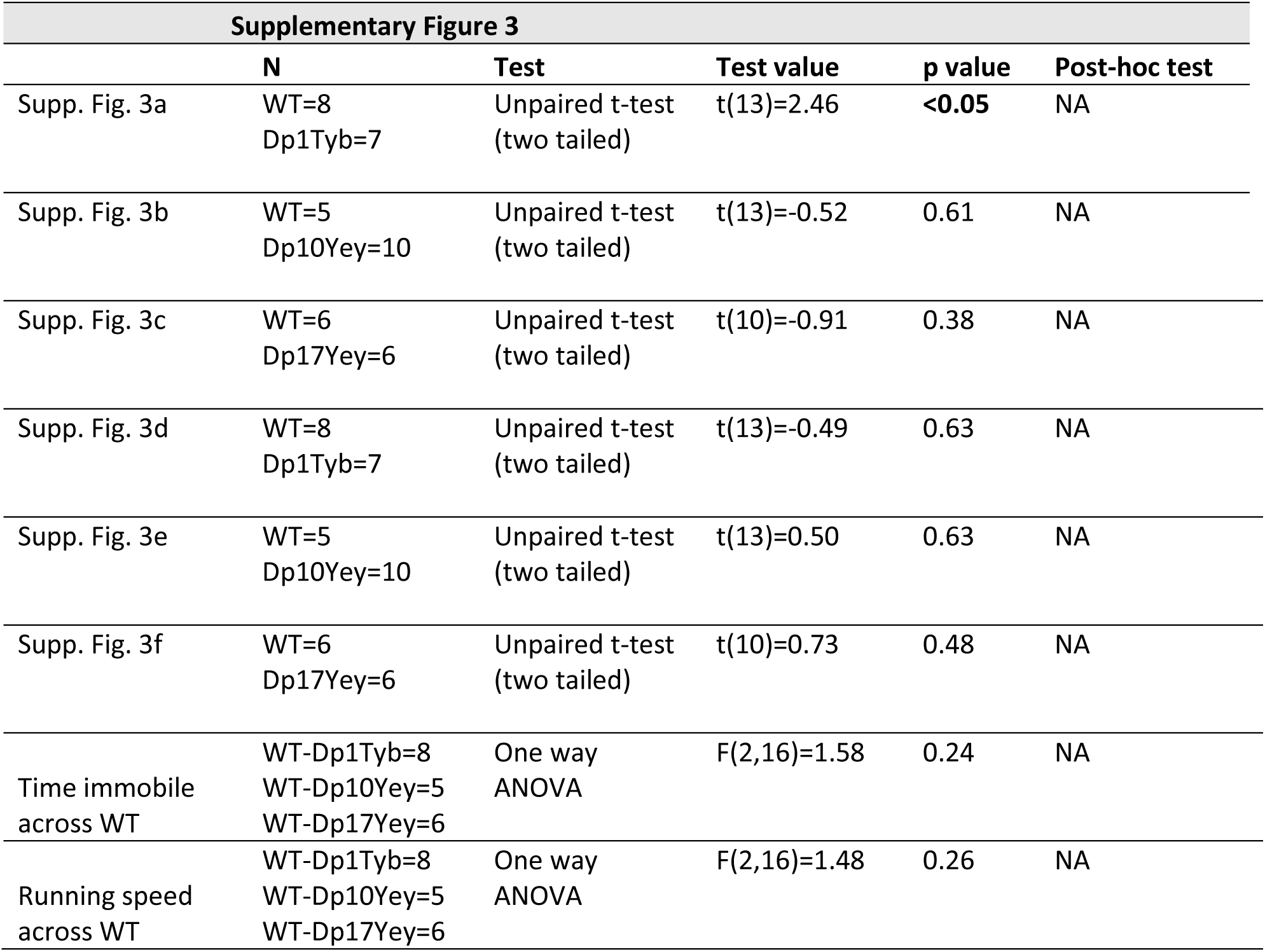
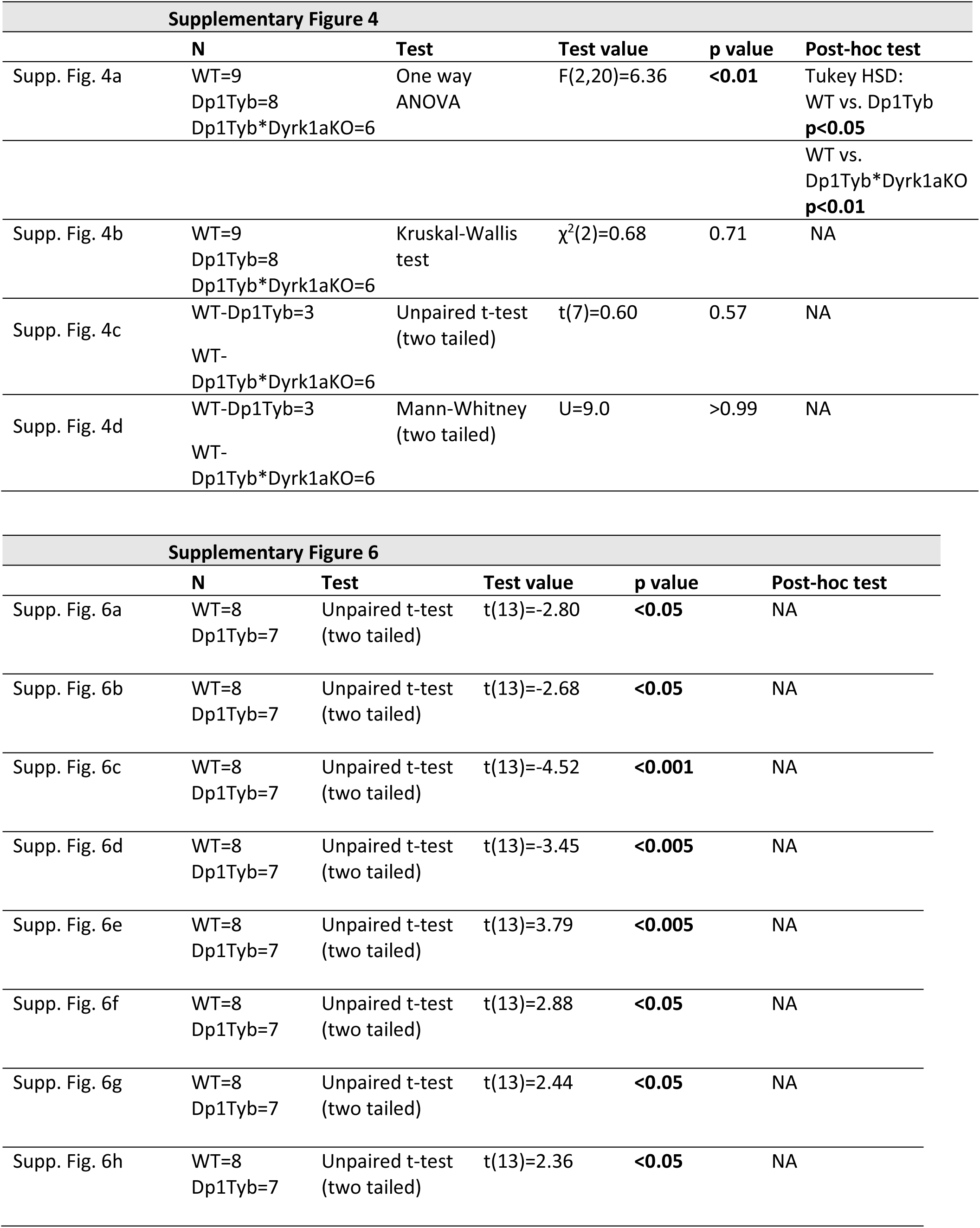
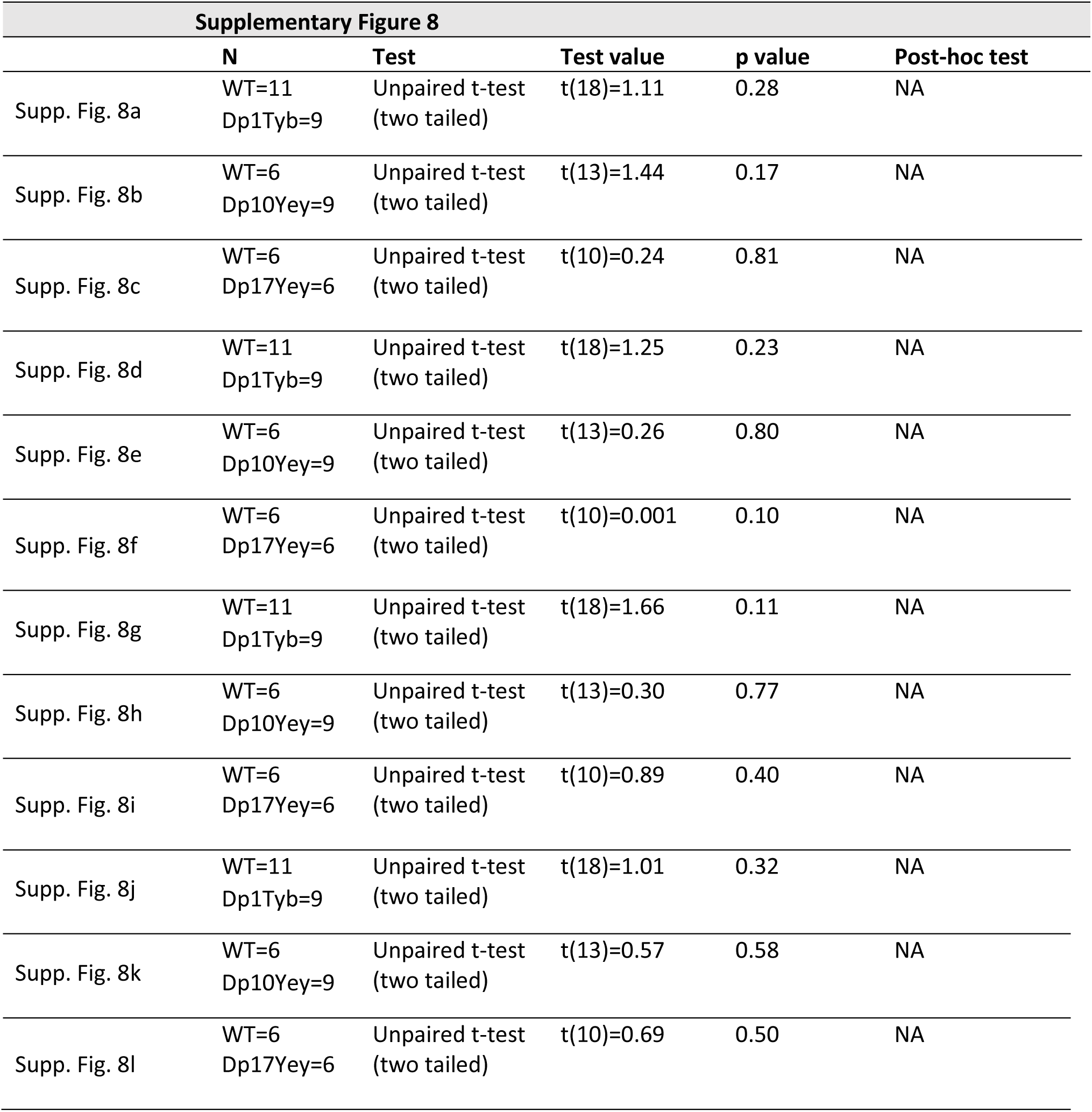
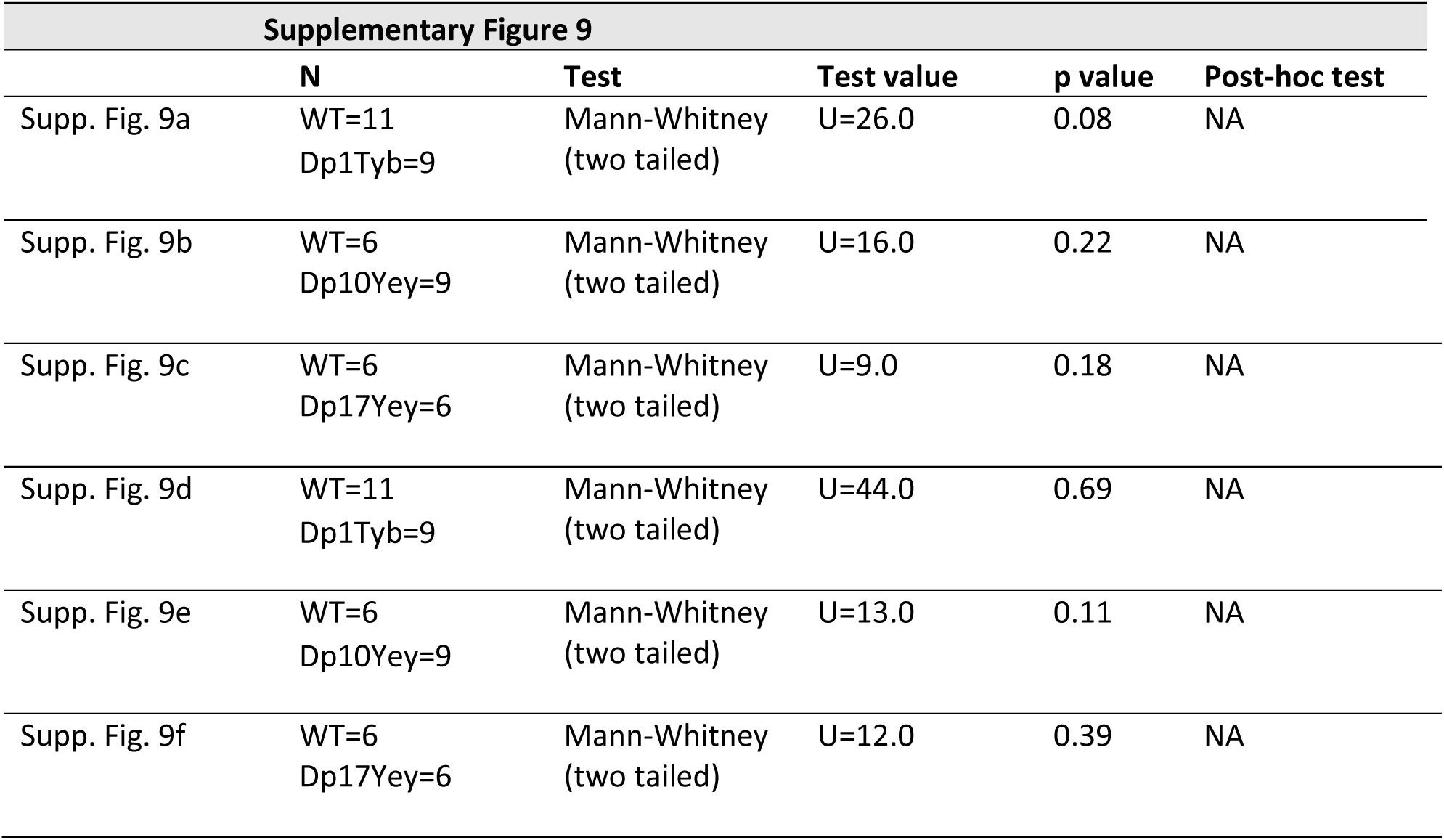
Full Details of all Statistical Analyses

**Supplementary Table 3:** Summary of Trial and Animal numbers for the Longitudinal Study

| 3 months |  |  |  |  |  |  |  |
| --- | --- | --- | --- | --- | --- | --- | --- |
|  | Dp1Tyb | WT | Dp10Yey | WT | Dp17Yey | WT | Mean |
| n | 4 | 6 | 5 | 3 | 6 | 5 | 4.83 |
| All Trials (Mean) | 8.75 | 8.17 | 9.40 | 6.67 | 10.17 | 9.80 | 8.83 |
| All Trials (SD) | 1.89 | 2.48 | 0.89 | 3.51 | 0.41 | 0.45 | 1.61 |
| Artefact Trials (Mean) | 2.50 | 1.17 | 2.00 | 0.00 | 1.00 | 0.40 | 1.18 |
| Artefact Trials (SD) | 3.32 | 2.40 | 3.39 | 0.00 | 0.89 | 0.89 | 1.82 |
| Included Trials (Mean) | 6.25 | 7.00 | 7.40 | 6.67 | 9.17 | 9.40 | 7.65 |
| Included Trials (SD) | 2.87 | 2.61 | 4.22 | 3.51 | 0.98 | 0.89 | 2.51 |

| 6 months |  |  |  |  |  |  |  |
| --- | --- | --- | --- | --- | --- | --- | --- |
|  | Dp1Tyb | WT | Dp10Yey | WT | Dp17Yey | WT | Mean |
| n | 4 | 6 | 5 | 3 | 6 | 5 | 4.83 |
| All Trials (Mean) | 12.50 | 10.83 | 10.00 | 10.33 | 10.00 | 10.00 | 10.61 |
| All Trials (SD) | 4.36 | 1.17 | 0.00 | 0.58 | 0.00 | 0.00 | 1.02 |
| Artefact Trials (Mean) | 0.00 | 0.33 | 0.00 | 0.00 | 0.67 | 0.20 | 0.20 |
| Artefact Trials (SD) | 0.00 | 0.52 | 0.00 | 0.00 | 0.82 | 0.45 | 0.30 |
| Included Trials (Mean) | 12.50 | 10.50 | 10.00 | 10.33 | 9.33 | 9.80 | 10.41 |
| Included Trials (SD) | 4.36 | 1.52 | 0.00 | 0.58 | 0.82 | 0.45 | 1.29 |

| 9 months |  |  |  |  |  |  |  |
| --- | --- | --- | --- | --- | --- | --- | --- |
|  | Dp1Tyb | WT | Dp10Yey | WT | Dp17Yey | WT | Mean |
| n | 3 | 4 | 4 | 2 | 6 | 5 | 4 |
| All Trials (Mean) | 10.33 | 12.25 | 10.25 | 10.00 | 10.17 | 10.40 | 10.57 |
| All Trials (SD) | 0.58 | 5.19 | 2.06 | 0.00 | 0.41 | 0.55 | 1.46 |
| Artefact Trials (Mean) | 0.00 | 0.25 | 0.00 | 0.50 | 0.33 | 0.00 | 0.18 |
| Artefact Trials (SD) | 0.00 | 0.50 | 0.00 | 0.71 | 0.52 | 0.00 | 0.29 |
| Included Trials (Mean) | 10.33 | 12.00 | 10.25 | 9.50 | 9.83 | 10.40 | 10.39 |
| Included Trials (SD) | 0.58 | 4.69 | 2.06 | 0.71 | 0.75 | 0.55 | 1.56 |

